# Epigenetic remodeling and 3D chromatin reorganization governed by NKX2-1 drive neuroendocrine prostate cancer

**DOI:** 10.1101/2024.12.04.626816

**Authors:** Xiaodong Lu, Viriya Keo, Irina Cheng, Wanqing Xie, Galina Gritsina, Juan Wang, Qiushi Jin, Peng Jin, Feng Yue, Martin G. Sanda, Victor Corces, Nicolas Altemose, Jonathan C. Zhao, Jindan Yu

## Abstract

A significant number of castration-resistant prostate cancer (CRPC) evolve into a neuroendocrine (NE) subtype termed NEPC, leading to resistance to androgen receptor (AR) pathway inhibitors and poor clinical outcomes. Through Hi-C analyses of a panel of patient-derived xenograft tumors, here we report drastically different 3D chromatin architectures between NEPC and CRPC samples. Such chromatin re-organization was faithfully recapitulated in vitro on isogenic cells undergoing NE transformation (NET). Mechanistically, neural transcription factor (TF) NKX2-1 is selectively and highly expressed in NEPC tumors and is indispensable for NET across various models. NKX2-1 preferentially binds to gene promoters, but it interacts with chromatin-pioneering factors such as FOXA2 at enhancer elements through chromatin looping, further strengthening FOXA2 binding at NE enhancers. Conversely, FOXA2 mediates regional DNA demethylation, attributing to NE enhancer priming and inducing NKX2-1 expression, forming a feed-forward loop. Single-cell multiome analyses of isogenic cells over time-course NET cells identify individual cells amid luminal-to-NE transformation, exhibiting intermediate epigenetic and transcriptome states. Lastly, NKX2-1/FOXA2 interacts with, and recruits CBP/p300 proteins to activate NE enhancers, and pharmacological inhibitors of CBP/p300 effectively blunted NE gene expression and abolished NEPC tumor growth. Thus, our study reports a hierarchical network of TFs governed by NKX2-1 in regulating the 2D and 3D chromatin re-organization during NET and uncovers a promising therapeutic approach to eradicate NEPC.

## INTRODUCTION

Following extensive treatment with androgen receptor (AR) pathway inhibitors, a significant number of metastatic prostate cancer develop treatment resistance, and approximately 20% of these castration-resistant prostate cancer (CRPC) transdifferentiate, at least partially, into neuroendocrine (NE) prostate cancer (NEPC) [1-4]. There are very limited treatment regimens for NEPC, and thus, understanding the molecular bases of NEPC is essential. At the genetic level, loss and aberrations of tumor-suppressor genes, such as PTEN, P53, and RB1, are frequently observed in NEPC and have shown a causative role [5-9]. Furthermore, dysregulated expression of lineage-specific transcription factors (TF), including MYCN, EZH2, and SOX2, have been shown to promote NE transformation (NET), primarily via suppressing AR signaling and yielding lineage plasticity [10-21]. Previous studies have also revealed substantial epigenetic changes in DNA methylation [3], chromatin accessibility [22], and histone modifications [23] in NEPC tumors. However, how these critical events that define cellular identity are programmed as PCa adenocarcinoma transdifferentiates into NEPC remains unknown.

Recent advances in the study of 3D chromatin folding have suggested that the human genome is hierarchically organized into multiple layers, including A/B compartments [24], topologically associating domains (TADs) [25], and chromatin loops [26]. The A and B compartments, roughly 1–10 Mb in length, are respectively associated with active and inactive chromatin. TADs and chromatin loops, approximately 0.1–1 Mb in size, are genomic regions that restrain or promote interactions between gene regulatory elements, such as enhancer-promoter (E-P) loops. Increasing evidence suggests that 3D chromatin structure, orchestrated by lineage-specific TFs, is cell type-specific and critical for cell fate determination [27-30]. However, although several studies have reported 2D epigenetic changes in NEPC [3], which will inevitably lead to alterations in the 3D chromatin architecture of a tumor, whether and how the 3D chromatin might be re-organized have not been investigated.

NKX2-1 (originally called TTF-1 for Thyroid transcription factor-1) is a homeodomain TF that is detected early in the endodermal thyroid and lung and in restricted neuroblast cells of the brain and is critical for their proper morphogenesis, acting as a lineage-specific TF [31-33]. NKX2-1 regulates the identity of neuronal progenitor cells, mediates interneuron specification, and directs postmitotic neuron migration [34, 35]. In the forebrain, NKX2-1 interacts with LHX6 to promote chromatin accessibility and activate genes expressed in the cortical migrating interneurons [36]. In the lung, NKX2-1 protein exhibits a temporal-spatial distribution similar to that of chromatin-pioneering factor FOXA2 in the developing and regenerating lung [37]. FOXA2 activates NKX2-1 transcription to regulate respiratory epithelial gene expression and cell differentiation [38]. NKX2-1, in turn, recruits FOXA proteins to lung-specific loci to promote growth and cellular identity while repressing gastric differentiation, acting as a master regulator favoring pulmonary differentiation [39-41]. In the prostate, NKX2-1 expression is undetectable in the luminal epithelium but is highly positive in more than 50% of NE lesions [42]. However, the roles of NKX2-1 in PCa have not been carefully studied.

Here, we mapped and revealed remarkably different 3D chromatin architecture in NEPC vs. CRPC tumors by Hi-C experiments. We recapitulated this 3D chromatin reorganization as isogenic prostate adenocarcinoma cells undergo NET in vitro and captured individual transitioning cells with intermediate chromatin and transcriptome states via scMultiome analyses. Mechanistically, we identify NKX2-1, a neural TF that is highly expressed in clinical NEPC samples, as a critical regulator of NE-lineage transdifferentiation in various NET models, while chromatin-pioneering factors such as FOXA2 prepare the chromatin through mediating regional DNA demethylation of target enhancers. Promoter-bound NKX2-1 interacts with enhancer-bound FOXA2 through E-P looping to facilitate FOXA2 binding at NE enhancers and induce NE enhancer activation through recruiting CBP/p300, ultimately leading to massive 3D chromatin reorganization and transcriptional reprogramming. Accordingly, CBP/p300 inhibition abolished NEPC growth *in vitro* and *in vivo*. Taken together, our study deciphers a general mechanism fundamental to cellular identity switch during NET and identifies potential therapeutic approaches using CBP/p300 inhibitors to deactivate NE enhancers and suppress NE tumor growth.

## RESULTS

### Distinct 3D chromatin architecture in NEPC compared to CRPC tumors

To examine the 3D chromatin architecture of PCa cells, we performed in situ Hi-C analyses of a panel of Patient-Derived Xenograft (PDX) tumors, including 4 CRPC and 3 NEPC, and 1 NEPC cell line NCI-H660 **(Supplementary Table 1**). We generated over 600 million valid paired-end reads for each sample, which were processed and summarized into a Hi-C interaction frequency matrix. Principal Component Analysis (PCA) was applied to each Hi-C matrix at a 100kb-sliding window to identify the eigenvector values of the first Principal Component (PC1), which are known to predict A/B compartments. To investigate whether NEPC contains unique chromatin compartmentalization, we performed unsupervised hierarchical clustering of all Hi-C samples based on their PC1 matrices. We found that CRPC and NEPC samples, except LuCaP147, were accurately separated into two distinct groups (**Fig.S1A**), suggesting significant compartmental differences between CRPC and NEPC tumors. Next, to further examine alterations in chromatin interactions, we identified chromatin loops at 10kb resolution using Mustache, p<0.05 [43]. Chromatin loops preferentially (p<0.001) detected in at least two NEPC samples in pairwise comparisons to CRPC samples were defined as NEPC-specific loops, and vice versa for CRPC-specific loops. As such, we identified 3,702 NEPC-specific loops and 1,808 CRPC-specific loops. Concordantly, Aggregate Peak Analysis (APA) of NEPC-specific loops demonstrated significantly higher APA scores in NEPC samples compared to CRPC samples, while the APA scores for CRPC-specific loops were substantially higher in CRPC samples (**Fig.1A-B**).

**Figure 1.**
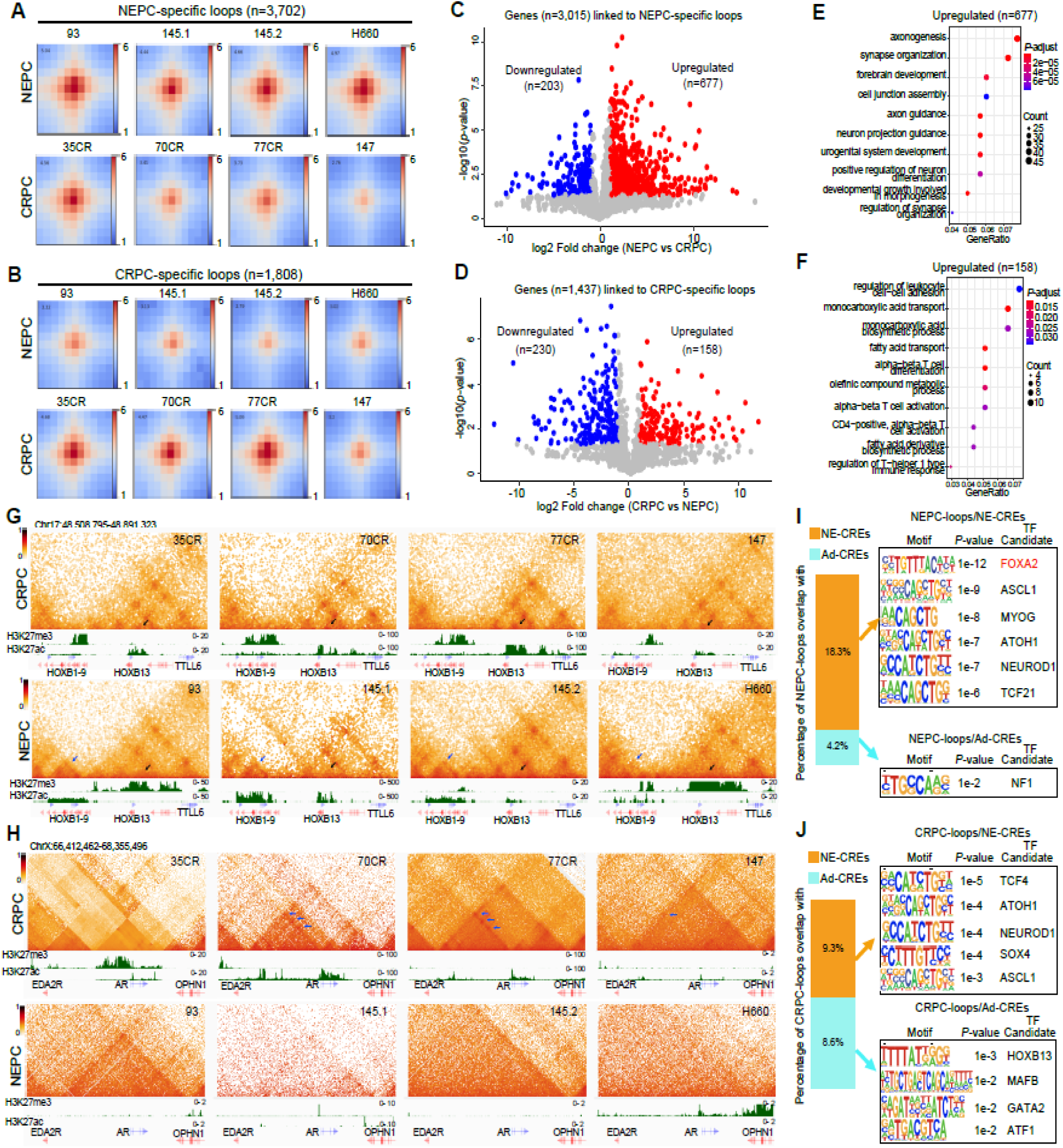
Distinct 3D chromatin architecture in NEPC compared to CRPC tumors. **A-B**. Aggregate Peak Analyses (APA) plots of NEPC-specific (**A**) and CRPC-specific (**B**) loops, respectively, in NEPC and CRPC samples. NEPC PDX: LuCaP93, LuCaP145.1, LuCaP145.2 and cell line: NCI-H660. CRPC PDX: LuCaP35CR, LuCaP70CR, LuCaP77CR, LuCaP147. **C-D**. Volcano plots showing genes linked to NEPC-(**C**) and CRPC-specific (**D**) loops that are differentially expressed (red/blue dots) in NEPC vs. CRPC samples, log2FC ≥ 1 and *P* < 0.05 by paired *t*-test. Public RNA-seq data of matched PDX (Labrecque, Coleman et al., Nouruzi, Ganguli et al.) was used for analyses. **E-F.** GO of genes linked to NEPC-(**E**) and CRPC-specific (**F**) loops and are up-regulated in NEPC and CRPC, respectively. **G-H.** Hi-C contact maps (5kb resolution) of the HOXB gene cluster (**G**) and AR gene region (**H)** in CRPC (**top row**) and NEPC (**bottom row**) tumors. Gene boxes and 1D ChIP-seq tracks of matched samples are aligned at the bottom. Arrows indicate NEPC/CRPC-specific loops. **I**-**J**. Bar graph showing the percentage of NEPC-(**I**) and CRPC-specific (**J**) loop anchors that overlap with previously reported (Baca, Takeda et al. 2021) NE-CREs (n=14,985) and Ad-CREs (n=4,338). The enriched motifs at overlapped anchors are shown on the right.

Chromatin looping is a significant mechanism for enhancer-mediated regulation of lineage-specific gene expression [44]. To assess potential targets of NEPC/CRPC-specific loops, we identified genes within ±3kb of loop anchors and examined their expression in public RNA-seq data of matched PDX [14, 45]. Interestingly, analyses of genes falling within NEPC-specific loop anchors revealed substantially more genes (n=677) that were significantly upregulated than down-regulated (n=203) in NEPC compared with CRPC (**Fig.1C**). By contrast, 230 and 158 genes associated with CRPC-specific loops were respectively down- and up-regulated in CRPC (**Fig.1D**), which may be due to chromatin loops functioning both as transcriptional activators and repressors as recently reported [44]. Gene Ontology (GO) analyses revealed that NEPC-loop genes that are up-regulated in NEPC are expectedly enriched in neuron development processes, such as axonogenesis and neuron projection guidance (**Fig.1E**). Consistently, CRPC-loop genes that are up-regulated in CRPC enriched for prostate functions, such as fatty acid transport, biosynthesis, and cell-cell adhesion (**Fig.1F**).

Next, we examined Hi-C contact maps at several representative NEPC- and CRPC-specific genes. For instance, we found a unique TAD at the HOXB1-9 gene cluster in NEPC that is not present in CRPC tumors (**Fig.1G**). Interestingly, this region appears to be epigenetically silenced by H3K27me3 in CRPC, which was erased in NEPC with a striking gain of active histone marker H3K27ac. Accordingly, RNA-seq analyses revealed that the expression of genes in the cluster, such as HOXB2, 7, and 9, were nearly undetectable in CRPC tumors but were abundantly expressed in NEPC (**Fig.S1B**). By contrast, epithelial TF gene HOXB13, located immediately next to the HOXB1-9 gene cluster, belonged to a different TAD that shows strong chromatin interactions in CRPC tumors, which were diminished in NEPC, with a concordant loss of H3K27ac and gain of H3K27me3 (**Fig.1G**). Such chromatin rewiring is consistent with recent reports of HOXB13 downregulation in NEPC via epigenetic silencing [46]. Moreover, a super-enhancer located ∼600kb upstream of the AR promoter has been shown to be critical for aberrant AR expression in CRPC [47]. Remarkably, we detected a strong chromatin loop between this enhancer and AR promoter, indicated by a unique TAD in CRPC tumors that was obliterated in NEPC (**Fig.1H and Fig.S1C**). In contrast with these lost TAD at luminal TFs in NEPC tumors, we observed gained Hi-C chromatin interactions at well-known NE regulators, such as ASCL1 and INSM1, with concordant epigenetic remodeling and gene activation (**Fig.S1D-G**).

Next, we attempted to exploit TFs that may be involved in orchestrating CRPC- and NEPC-specific chromatin looping. To focus on enhancer loops, we overlapped previously defined NE-enriched Candidate Regulatory Elements (NE-CREs) and adenocarcinoma-enriched CREs (Ad-CREs) [23] with loop anchors. Our results showed that remarkably more NEPC-specific loops overlapped with NE-CREs (18.3%) than Ad-CREs (only 4.2%), and they are strongly enriched for the DNA binding motif of FOXA2, a biomarker gene of NEPC [48, 49] that has been shown to play a causative role (**Fig.1I**) [50, 51]. By contrast, CRPC-specific loops overlapped similarly with NE- and Ad-CREs, which were enriched for motifs of NE and luminal TFs, respectively (**Fig.1J**). Altogether, our data demonstrated striking differences in the 3D chromatin architecture between NEPC and CRPC tumors with disease-specific chromatin interactions that are associated with lineage-specific enhancer activation and gene expression.

### Remarkable 3D chromatin reorganization during NE transformation of isogenic PCa cells

We wondered whether the substantial differences in the 3D chromatin architecture in NEPC vs. CRPC tumors can be driven by individual NE regulators. As FOXA2 has been shown to promote NET of prostate cancer [50, 51] and its motif was enriched in our NEPC-specific Hi-C loops, we infected LNCaP (adenocarcinoma) cells with FOXA2 lentivirus and monitored the cells over a time course of 28 days. Interestingly, approximately 30% of FOXA2-overexpressing cells (LNCaP+FOXA2) exhibited NE phenotype with neurite extensions and formed “cluster” morphology at day 21 (D21), which increased to over 90% at D28 (**Fig.S2A**). In agreement with this, western blot (WB) analyses revealed gradually reduced expression of luminal TFs AR and HOXB13 over time and increased expression of NE lineage markers, such as SYP, NCAM1, and BRN2, manifesting at D21 and peaking at D28 (**Fig.S2B**). Similarly, FOXA2 overexpression (OE) led to NET of 22Rv1 cells over a period of 28 days and of the AR-negative DU145 cells in only 7 days (**Fig.S2C**). Next, to examine global differential gene expression, we performed RNA-seq analyses of time-course LNCaP+FOXA2 cells. We found that a large number of genes (n=12,835, DESeq2 likelihood ratio test, adjust p<0.0001) were differentially expressed, indicating major transcriptional reprogramming. K-means clustering revealed 6 clusters, including 3 clusters each of up-regulated (C1-C3, n=6,825) and down-regulated genes (C4-6, n=6,010) (**Fig.2A**). GO analyses showed that the biggest gene clusters, C2 and C5, respectively induced gene involved in neurogenesis and embryonic development (e.g., SOX2 and POU3F2) and repressed genes essential for epithelial functions (e.g., HOXB13 and AR), with the molecular switch occurring between D14 and D21. Early-response clusters of genes (C1 and C4) increased DNA damage response and decreased rRNA processing that occurred between D2 and D7, indicating cellular stress. Interestingly, the cells might have coped with cellular stress by transiently reducing mRNA metabolism (C3) and increasing lipid biosynthesis (C6) until they reach a stable NE-transformed state. Furthermore, to examine how well our time-course data recapitulate the transcriptome of patient-derived samples, we performed integrative analyses [52] of AR and NE signature scores in our RNA-seq data and that of clinical PCa samples from a previous study [45]. Critically, D0, D2, and D7 cells all clustered with AR+/NE- tumors, as expected, while D21 and D28 cells were more closely grouped with AR-/NE+ (NEPC) tumors (**Fig.2B**). Importantly, D14 cells were somewhere in between, with intermediate AR and NE signature expression. This data suggests that our time-course cells faithfully modeled the clinical transition of PCa from adenocarcinoma to NEPC and identified D14 as the tipping point of the transition.

**Figure 2.**
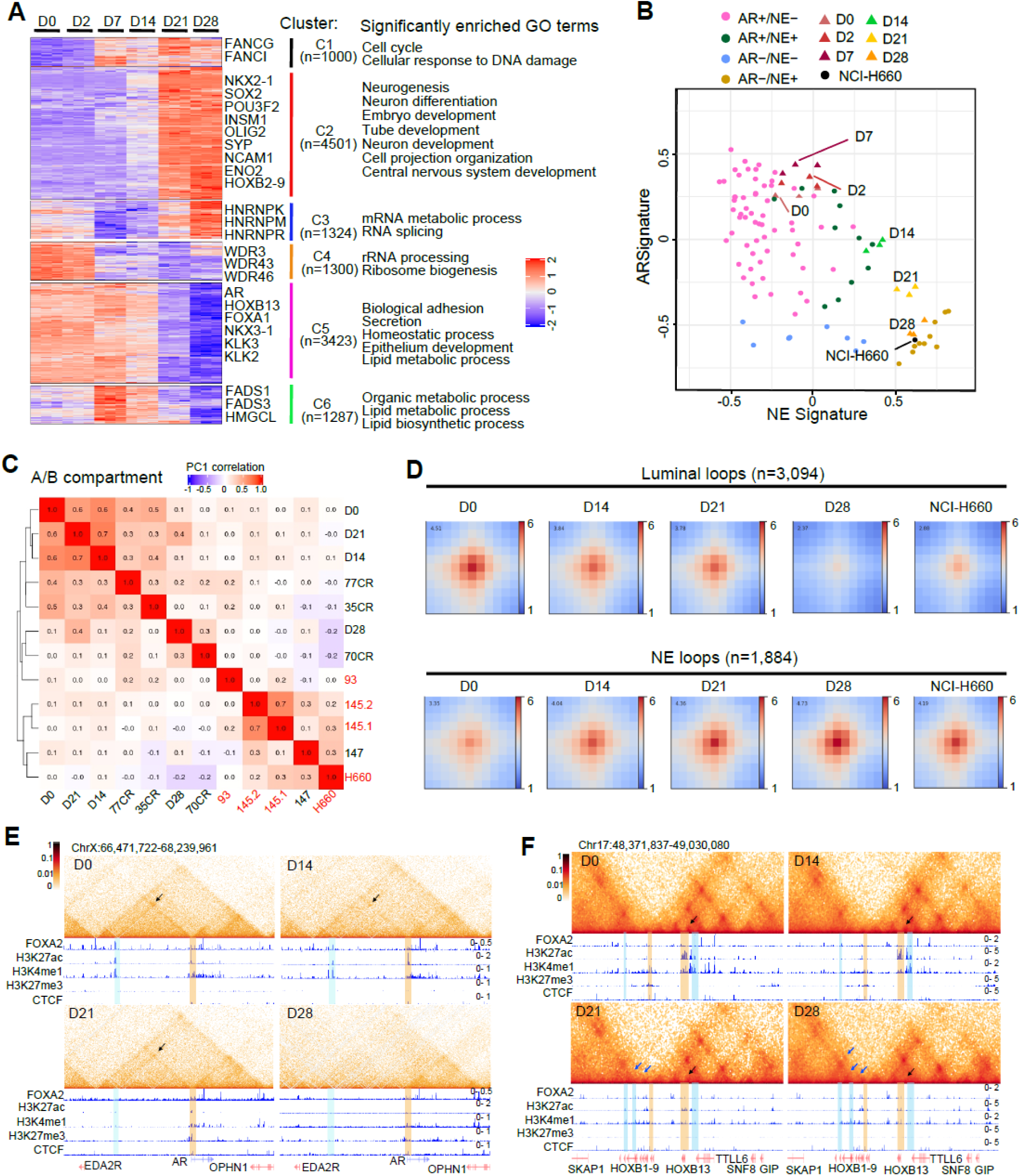
Remarkable 3D chromatin reorganization during NE transformation of isogenic PCa cells. **A**. Heatmap shows 6 clusters of genes differentially expressed over time course (adjusted p<0.0001). Representative genes and significantly enriched molecular concepts are shown on the right. Bulk RNA-seq was performed in LNCaP cells with FOXA2 OE at indicated time points. **B**. Integrative analyses of AR and NE signature scores in time-course LNCaP+FOXA2 samples and that of clinical PCa samples (GSE126078). AR and NE signature scores were calculated by Gene Set Variation Analysis (GSVA). **C**. Unsupervised hierarchical clustering of NEPC, CRPC, and time-course LNCaP+FOXA2 samples based on the top 10% most variable PC1 of Hi-C matrices. **D**. APA plots of luminal (**top row**) and NE (**bottom row**) loops (D28 vs. D0) in time-course LNCaP+FOXA2 samples and NCI-H660. **E-F**. Hi-C contact maps (5kb resolution) of AR region (**E**) and HOXB cluster (**F**) in time-course LNCaP+FOXA2 cells. Gene boxes and 1D ChIP-seq tracks of matched cells are aligned at the bottom. Black arrows indicate luminal loops at AR and HOXB13 locus, and blue arrows indicate NE loops at HOXB1-9 loci. Enhancers are highlighted by light-blue columns and promoters light orange. FOXA2, H3K27ac, H3K4me1, H3K27me3, and CTCF tracks under the D0 contact map utilized their corresponding ChIP-seq performed in closely related D2 cells. All others were done in conditions that exactly matched Hi-C.

To explore whether the 3D chromatin organization is rewired during FOXA2-induced NET, we performed in situ Hi-C of time-course LNCaP+FOXA2 cells. First, hierarchical clustering of Hi-C data using their PC1 matrices demonstrated a tight clustering of D0, D14, and D21 samples with CRPC tumors, whereas D28 cells were loosely clustered with NEPC and some CRPC tumors (**Fig.2C**). This data indicates a large-scale chromatin compartmental re-organization following FOXA2 OE that recapitulates clinical CRPC transformation to NEPC. Furthermore, to examine the chromatin architecture during NET at the loop level, we identified 3,094 luminal loops and 1,884 NE loops by comparing D0 with D28. As expected, the NE loops potentially regulate genes implicated in axonogenesis and developmental processes, and the luminal loops control genes involved in epithelial functions (**Fig.S2D**). APA analyses demonstrated that the luminal loops were gradually attenuated from D0 to D28, while the NE loops were strengthened over time (**Fig.2D**). Interestingly, the APA scores of D28 cells were comparable to those of NCI-H660, supporting a re-organized 3D chromatin architecture that is similar to prototype NEPC cells. For example, we detected the loop between the AR promoter and the ∼650kb upstream enhancer in D0 cells, being consistent with our findings in CRPC PDX tumors (**Fig.2E**). Critically, this E-P loop was gradually lost over the time-course. Accordingly, there was a decrease of H3K4me1 and H3K27ac, histone marks for enhancer priming and activation, respectively, from D0 to D28. By contrast, broad peaks of H3K27me3 were detected at the AR gene, starting at D21, suggesting concomitant epigenetic remodeling along with 3D chromatin rewiring. Likewise, we observed contrasting changes in epigenetic modifications and TADs at the HOXB1-9 cluster and HOXB13 gene over the time course, which is strikingly similar to what we have seen when comparing CRPC to NEPC PDX tumors (**Fig.2F**). Therefore, our data demonstrate that the distinct 3D chromatin architecture observed in patient-derived CRPC and NEPC tumors can be recapitulated in isogenic cells transforming from luminal to NE lineage. Our LNCaP+FOXA2 system thus represents a clinically relevant model for deciphering the molecular mechanisms and kinetics underlying NET.

### Single-cell multiome analyses identified individual transitioning cells with intermediate transcriptome and chromatin states

To understand the epigenetic bases of the transcriptional reprogramming and 3D chromatin re-organization that we have observed during NET, we performed Assay for Transposase-Accessible Chromatin using sequencing (ATAC-seq) in time-course LNCaP+FOXA2 cells and identified 89,636 chromatin regions with significantly (adjusted *p*-value<0.0001) altered accessibility. K-means clustering of ATAC-seq peaks revealed 6 major clusters, including 2 clusters (C1-C2) of peaks that are shared over all time points, 2 clusters (C3-C4) of gradually increasing peaks, and 2 clusters (C5-C6) of gradually decreasing peaks (**Fig.3A and Fig.S3A**). The C3-C4 peaks became detectable only at D14 and localized primarily at enhancer elements, hereafter defined as NE enhancers (**Fig.3B**). Motif analyses indeed identified FOXA2, NKX2-1, and developmental and stem cell TFs, such as SIX1/2/4, SOX2, and OCT4, as the top enriched TFs at these regions (**Fig.3A**). Furthermore, GO analyses revealed that the genes within 50kb of these peaks were involved in neuron fate specification and stem cell development/function. On the contrary, C5-C6 peaks were strong in D0-D7 luminal cells and became inaccessible over time, thus defined as luminal enhancers (**Fig.3B**). These peaks were enriched for luminal TFs such as FOXA1, AR (ARE motif), and HOXB13 and correspond to genes involved in prostate functions. The shared peaks, especially C1, were predominantly at promoter regions and enriched for motifs of basic TFs, such as SP1, likely critical for housekeeping gene expression. A comparison of enriched motifs in D2 vs. D28 ATAC-seq peaks revealed NKX (including NKX2-1/2/5), OCT4, and ISL1 motifs as top ranked in D28 cells, whereas motifs of prostate-specific/luminal TF FOXA1, GATA3 and AR (ARE motif) ranked highly in D2 cells (**Fig.S3B**). PCA analyses using differential ATAC-seq peaks over LNCaP+FOXA2 time course [22] showed that D0/2/7 cells and D21/28 cells were respectively clustered closer to CRPC and NEPC PDX tumors, whereas the D14 cells were positioned somewhere in between, indicating a transitioning stage (**Fig.S3C**). Altogether, these results support an evident chromatin remodeling that is consistent with luminal-to-NE cellular switch from D2 to D28 cells and resembles clinical CRPC/NEPC samples.

**Figure 3.**
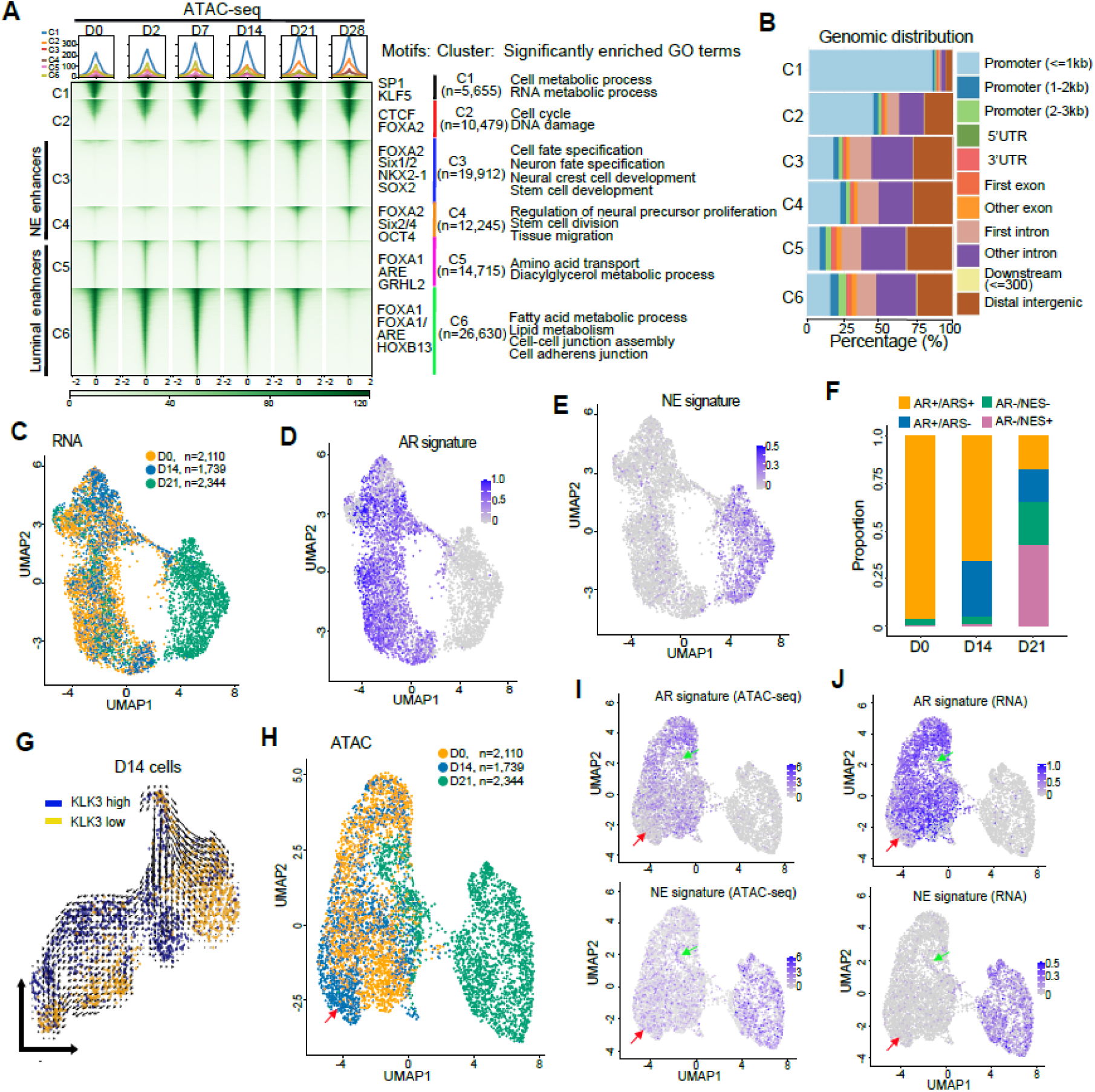
Single-cell multiome identified transitioning individual cells with intermediate transcriptome and chromatin states. **A**. K-mean clustering reveals 6 clusters of differential ATAC-seq peaks (±2kb) across time-course LNCaP+FOXA2 samples (adjusted p<0.001). Heatmaps shown here were scaled across samples. Color bar at the bottom indicates the scale of enrichment intensity. Significantly enriched TF motifs and GO terms for each cluster, along with the number of peaks, are shown on the right. **B**. Genomic distribution of the 6 clusters of ATAC-seq peaks in **A**. **C-F**. scRNA-seq UMAP visualization of D0, D14, and D21 LNCaP+FOXA2 cells (**C**), AR (**D**) and NE (**E**) signature genes, which are further quantified for the proportion of AR+/ARS+, AR+/ARS-, AR-/NES-, AR-/NES+ cells (**F**). **G**. RNA velocity analysis of D14 LNCaP+FOXA2 cells. Data were visualized as streamlines in a UMAP-based embedding. Blue dots indicate KLK3-high cells, and yellow dots KLK3-low cells. **H-J**. scATAC-seq UMAP visualization of D0, D14, and D21 LNCaP+FOXA2 cells (**H**), chromatin accessibility (**I**), and mRNA levels (**J**) of AR/NE signature genes. Red and green arrows indicate transitioning D14 and D21 cells, respectively.

To affirm that the observed epigenetic and transcriptional reprogramming is due to cell transformation, rather than populational switch, we exploited single-cell (sc) Multiome (scRNA-seq and scATAC-seq) to capture individual, transitioning cells with intermediate transcriptome and chromatin states. We captured 2,110 D0, 1,739 D14, and 2,344 D21 LNCaP+FOXA2 cells for scMultiome analyses. First, comparing the transcriptome status of our cells with individual clinical PCa cells [53] revealed that the majority of our D0 cells were clustered in close proximity to primary PCa cells, most of the D14 and some of the D21 cells were grouped along with CRPC cells, while a majority of D21 cells are in the neighborhood of NEPC cells (**Fig.S3D**). Trajectory analyses further validated that our D14 cells and patient CRPC cells were in the middle stage of the pseudotime (**Fig.S3E**). These data confirm the clinical relevance of our model. Next, we performed Uniform Manifold Approximation and Projection (UMAP) analyses of our scRNA-seq data, which revealed two distinct clusters that comprised primarily of D0 and D21 cells and expressed high levels of AR and NE genes, respectively (**Fig.3C & S3F**). Of note, most of the D14 and some of the D21 cells still belonged to the D0 cluster. These cells remained AR-positive, but had a reduced AR signature (ARS) score and started to express some NE signature (NES) genes (**Fig.3D-E**), indicating an intermediate transcriptome. To investigate this more closely, we quantified the proportion of cells at each time point that express AR, ARS, or NES genes and found that a majority (96.5%) of D0 cells are AR+/ARS+, as expected for androgen-dependent PCa (**Fig.3F**). Critically, this number dropped to around 55% at D14 with the emergence of a new cell type (∼30%) that is still AR+ but has lost ARS expression. AR itself was ultimately depleted in most (∼65%) of the D21 cells, a majority of which also gained NES expression (AR-/NES+). To provide further evidence that the AR+/ARS-D14 cells are a result of transcriptional reprogramming of the original AR+/ARS+ cells, we performed RNA velocity analyses of D14 cells. Indeed, we detected a strong tendency of the KLK3-high (AR+/ARS+) cells to become KLK3-low (AR+/ARS-) (**Fig.3G**). These data identify individual D14 and D21 cells that are in the midst of transition, with an intermediate transcriptome. Similarly, UMAP analyses of matched scATAC-seq data also revealed two distinct clusters: a pure cluster of most D21 cells and a mixed cluster of D0, D14, and some D21 cells (**Fig.3H**). Focusing on chromatin accessibility at the promoter and intragenic regions of AR and NE signature genes, we found that the chromatin of AR signature genes was accessible mostly in D0 cells, whereas the chromatin of NE signature genes was much more accessible in D21 cells, as expected (**Fig.3I**), with concordant changes in signature expression (**Fig.3J**). Importantly, although most D14 cells clustered together with D0 cells, many D14 cells showed inaccessible chromatin at AR signature genes and modest chromatin accessibility at NE genes (**Fig.3I, red arrows**), similar to the D21 cells within the cluster (**Fig.3I, green arrows**). In aggregates, the presence of individual D14 and D21 cells with matched, intermediate chromatin and transcriptome states strongly supports luminal cell transformation to a new NE lineage, leading to NET.

### NKX2-1 is induced by FOXA2 and required for FOXA2-driven NET

We have observed that FOXA2 OE in LNCaP cells took 28 days to complete NET, with D14 as the tipping point of transcriptomic and epigenetic reprogramming, which may be triggered by some driver events that only become available at D14. As FOXA2 is a pioneering factor, which requires the coordination of lineage-specific TF for function, we focused on neural TFs that are turned on at D14. We found that NKX2-1 was absent in D0 and D2, started to have some expression at D14, and reached full expression at D21 (**Fig.4A**). ChIP-seq data showed multiple FOXA2 binding events at upstream and downstream enhancers of NKX2-1 as early as D2, which reached a high level at D14, supporting FOXA2 binding as an initiating event inducing NKX2-1 expression. Indeed, H3K4me1 at the NKX2-1 enhancers became detectable at D14, suggesting it’s being primed by FOXA2 binding. H3K27ac lagged slightly until D21, indicating that additional efforts are required for the activation of primed enhancers. These sequential events strongly support that FOXA2 binds to NKX2-1 enhancers to initiate and induce its expression. FOXA2 occupancy at NKX2-1 enhancers were confirmed in additional NEPC models, including NCI-H660 and LuCaP145.2 (**Fig.S4A**). Moreover, FOXA2 knockdown (KD) in LuCaP145.2 organoids, NCI-H660, and stable LNCaP+FOXA2 cells (>D28), hereafter defined as <u>LuNE</u> cells, drastically reduced NKX2-1 expression (**Fig.4B**). Taken together, these data indicate that FOXA2 directly induces NKX2-1 gene transcription.

**Figure 4.**
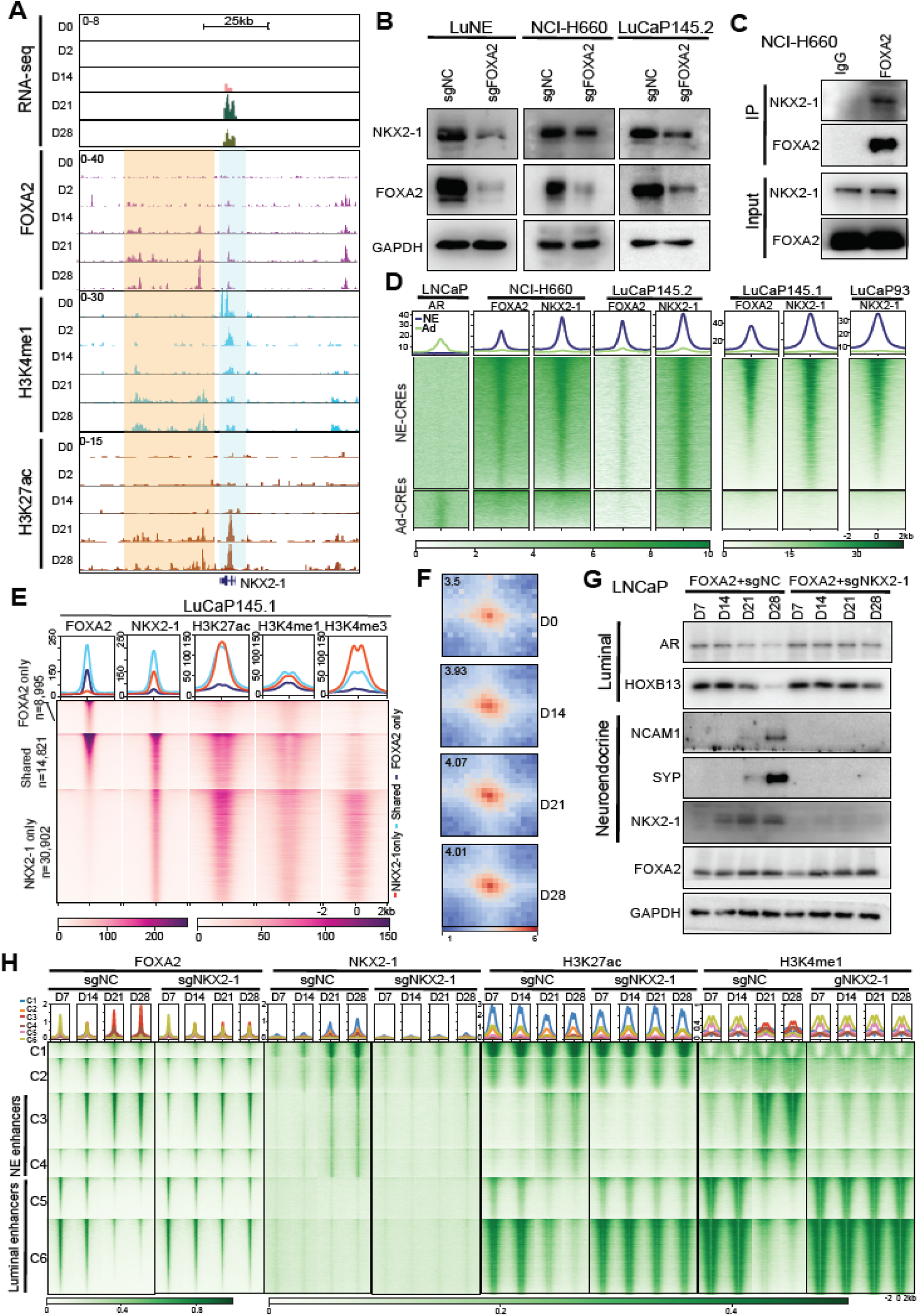
NKX2-1 is induced by FOXA2 and is required for FOXA2-driven NET. **A.** Genome browser view of RNA-seq and FOXA2, H3K4me1, and H3K27ac ChIP-seq signal around the NKX2-1 gene in time-course LNCaP+FOXA2 cells. The gene is highlighted in light blue, and enhancers in yellow box. **B.** WB of NEPC cell lines and organoids (LuCaP145.2) with control or FOXA2 KD. **C.** FOXA2 Co-IP showing its interaction with NKX2-1 protein in NEPC cell line NCI-H660. **D.** Heatmap showing AR, NKX2-1, and FOXA2 occupancy at previously reported (Baca, Takeda et al. 2021) NE-CREs (n=14,985) and Ad-CREs (n=4,338) in LNCaP or NEPC models (NCI-H660, LuCaP93, LuCap145.1, LuCaP145.2). **E.** Heatmap showing active (H3K27ac) enhancer (H3K4me1) and promoter (H3K4me3) marks at NKX2-1-only, FOXA2-only, and shared binding sites in LuCaP145.1. **F.** APA plots of FOXA2- and NKX2-1-anchored loops identified specifically in D28 LNCaP+FOXA2 Hi-C data. **G.** NKX2-1 KD abolishes FOXA2-induced NET. LNCaP cells were co-infected with FOXA2 virus along with either sgNC or sgNKX2-1 virus. The infected cells were collected at indicated time points for WB analyses of luminal and NE markers. **H.** Heatmap showing FOXA2, NKX2-1, H3K27ac, and H3K4me1 ChIP-seq around (±2kb) the 6 ATAC-seq peak clusters in LNCaP+FOXA2 cells, with or without NKX2-1 KD, at indicated time points. Peaks within each cluster were separately sorted by FOXA2 ChIP-seq intensity. Scale bar: enrichment intensity.

To determine whether NKX2-1 facilitates FOXA2 function, we first performed co-immunoprecipitation (co-IP) in NCI-H660, which showed strong interactions between endogenous FOXA2 and NKX2-1 proteins (**Fig.4C**). To evaluate whether they could be functionally involved in regulating NE enhancers, we conducted ChIP-seq in various NEPC models, which indeed revealed substantial NKX2-1 and FOXA2 co-occupancy at previously defined NE-CREs [23] (**Fig.4D**), while AR binds to only Ad-CREs. Further, we found that a majority of FOXA2-binding sites co-occupied by NKX2-1 were marked by strong H3K27ac and H3K4me1, suggesting active NE enhancers. In contrast, FOXA2-only binding sites were marked with H3K4me1 only, indicative of primed enhancers, whereas a significant number of NKX2-1-only binding sites are active promoters enriched for both H3K4me3 and H3K27ac (**Fig.4E and Fig.S4B-C**). Next, we attempted to understand how NKX2-1, which predominantly binds to promoters, facilitates and collaborates with the enhancer-bound FOXA2. Considering that NKX2-1 and FOXA2 proteins interact, we hypothesized that their interaction may be made possible by chromatin looping. Analyzing loops that are anchored at FOXA2 and NKX2-1 binding sites in LNCaP+FOXA2 time-course Hi-C data, we observed a substantial increase in the number of chromatin loops, from 5 at D0 to 1,305 at D28 (**Fig.S4D**). Moreover, APA analysis showed drastically increased intensity of chromatin loops that are anchored at FOXA2 and NKX2-1 binding sites (**Fig.4F**). These results suggest that FOXA2 and NKX2-1 proteins interact and co-occupy NE enhancers through chromatin looping.

Lastly, we sought to investigate whether the induction of NKX2-1 by FOXA2 at D14 is required to tip the balance to a complete NET. To this end, we performed FOXA2 OE in LNCaP cells with concurrent control or NKX2-1 KD. Remarkably, the depletion of NKX2-1 entirely abolished FOXA2-driven NET, indicated by the lack of switches in luminal and NE marker gene expression in cells with FOXA2 OE and NKX2-1 KD (**Fig.4G**). To evaluate how epigenetic states are regulated by FOXA2 and NKX2-1 during FOXA2-induced NET, we carried out FOXA2, NKX2-1, H3K27ac, and H3K4me1 ChIP-seq in time-course LNCaP cells subjected to FOXA2 OE and concomitant control or NKX2-1 KD. Interestingly, immediately following OE, FOXA2 bound to luminal enhancers that are marked by H3K4me1 and H3K27ac, which is consistent with previous reports of such epigenetic signatures in recruiting FOXA pioneering factors [54] (**Fig.4H**). By contrast, NKX2-1 did not have detectable binding events at early points, which is consistent with it being expressed only after D14. Of note, once turned on at D21, NKX2-1 only binds to NE, but not luminal, enhancers. Importantly, there was a remarkable shift of FOXA2, H3K27ac, and H3K4me1 binding from luminal to NE enhancers at D21 and beyond, suggesting that NKX2-1 expression might be essential for epigenetic programming. In support of this, depletion of NKX2-1 expression completely abolished this lineage switch, as manifested by continued H3K27ac and H3K4me1 binding at luminal enhancers at D21 and D28 of NXK2-1-KD cells (**Fig.4H**). Further, there was a slight shift of FOXA2 cistrome from luminal to NE enhancers that was also halted at D14 in NKX2-1-KD cells. These results showed that FOXA2 initiates lineage plasticity but requires the induction of NKX2-1 to complete the luminal-to-NE switch, suggesting a critical role of NKX2-1 in NEPC.

### NKX2-1 is highly expressed in NEPC tumors and critical for NET in various models

NKX2-1 has been implicated in pulmonary differentiation, but its role in PCa has not been well studied. We analyzed the expression of all human TFs in 3 large publicly available PCa datasets [3, 45, 55] and found NKX2-1 and FOXA2 to be significantly co-elevated in NEPC, while AR was enriched in CRPC, as expected (**Fig.5A and S5A**). Further analyses showed remarkable up-regulation of NKX2-1 and FOXA2 mRNA in the AR-/NE+ (NEPC) subtype compared with AR+/NE-, AR+/NE+, and AR-/NE-tumors (**Fig.5B**). To confirm this at the protein level, we performed immunohistochemistry (IHC) staining in tissue microarrays (TMA) containing clinical PCa samples. We validated strongly elevated levels of NKX2-1 and FOXA2 proteins in AR-/NE+ samples relative to other subtypes of PCa (**Fig.5C**). Moreover, while FOXA2 was increased in some AR-/NE-as well as AR-/NE+ tumors, NKX2-1 was uniquely up-regulated in NEPC (AR-/NE+) tumors (**Fig.5B-C**), suggesting a critical role of NKX2-1 in specifically regulating NEPC.

**Figure 5.**
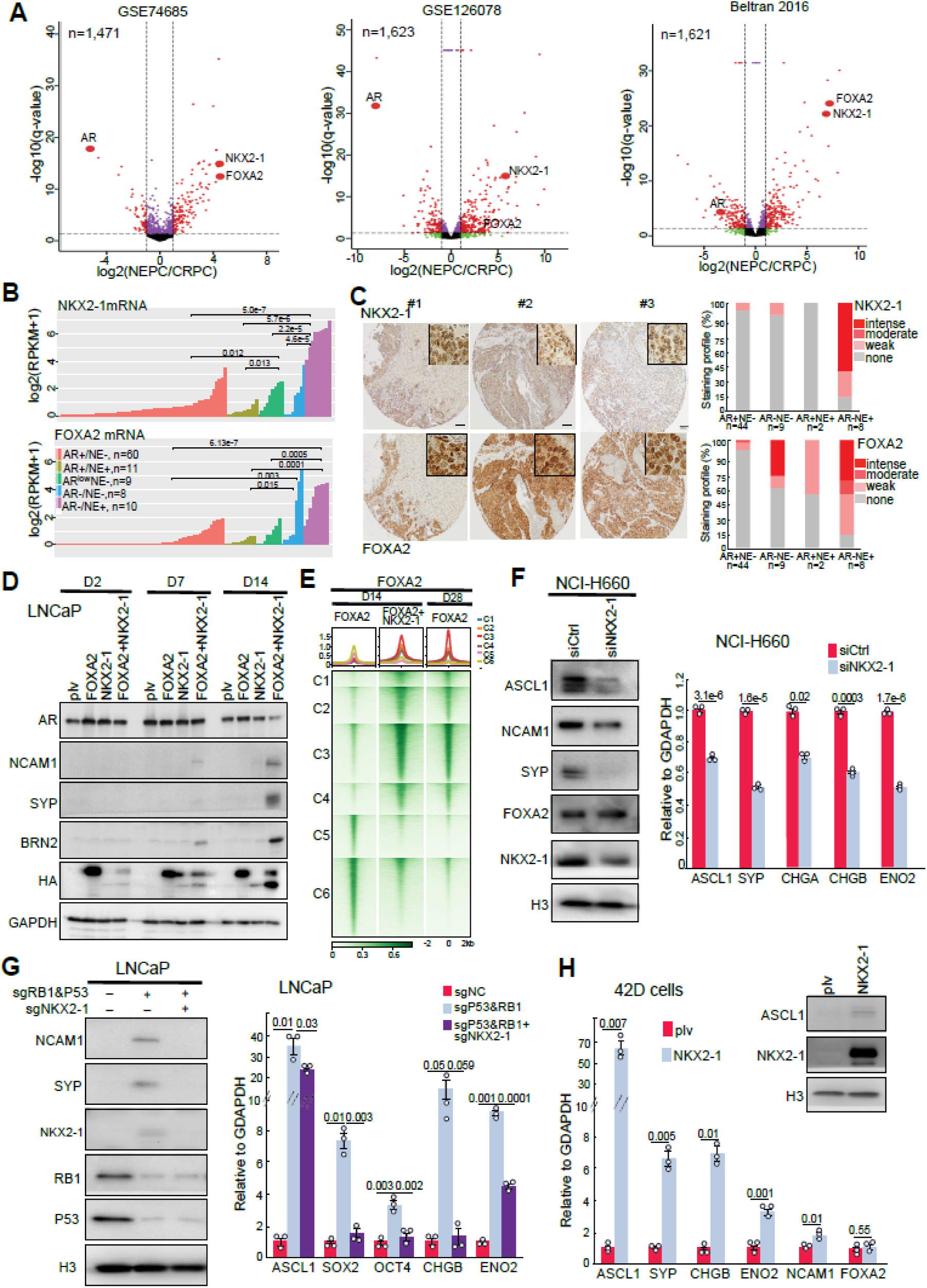
NKX2-1 is highly expressed in NEPC tumors and critical for NET in various models. **A.** Volcano plots showing human transcription factors (TFs) differentially expressed in CRPC vs. NEPC. X-axis represents log2 of fold change (FC), while y-axis shows −log10 of q values. Each dot represents a TF, red dots: log2FC ≥1 and adjusted *p* < 0.05. **B.** NKX2-1 and FOXA2 gene expression in human PCa samples (GSE126078) with distinct AR and NE gene expression. *P*-values by Wilcoxon test. **C.** IHC of NKX2-1 and FOXA2 proteins in clinical CRPC/NEPC TMAs. Representative IHC images (3 independent cores) are shown in 10x, with the inserts shown in 40x **(left**). IHC intensities in AR+NE-, AR-NE-, AR+NE+, and AR-NE+ samples were quantified on the right. **D.** Concomitant NKX2-1 OE accelerates FOXA2-driven NET. LNCaP cells were infected with FOXA2, NKX2-1, or both. Cells were collected at indicated time points and analyzed by WB. **E.** Heatmap showing FOXA2 ChIP-seq in LNCaP cells with overexpression of FOXA2 alone, or FOXA2 and NKX2-1 at D14 or D28. Peaks centered around (±2kb) the 6 ATAC-seq clusters and sorted by FOXA2 ChIP-seq intensity. Scale bar: enrichment intensity. **F.** NKX2-1 KD decreases the expression of NE lineage markers in NCI-H660. NCI-H660 cells were transfected with control or NKX2-1 siRNAs and harvested on day 5 after transfection for WB and RT-PCR analyses. RT-PCR data were normalized to GAPDH. Shown are the mean ± s.e.m. of technical replicates from one of three (n = 3) independent experiments. *P* values were calculated by unpaired two-sided *t*-test **G.** NKX2-1 KD abolishes RB1/P53 KD-induced NET. LNCaP cells were co-infected with sgRB1 and sgP53 virus along with either sgNC or sgNKX2-1 virus. The infected cells were collected at 4 weeks after infection for analyses of NE and stem cell lineage markers. **H.** NKX2-1 OE promotes NET in enzalutamide-resistant AR+/PSA− cell line 42D. The infected cells were collected at 4 weeks after infection for analyses of NE lineage markers.

To investigate the roles of NKX2-1 in driving NET, we overexpressed NKX2-1 in LNCaP cells, similar to FOXA2 OE. However, we failed to observe NET morphology and SYP marker expression after 4 weeks of NKX2-1 OE, despite some decrease in AR and PSA levels (**Fig.S5B**). We thus hypothesized that chromatin preparation by pioneering factors such as FOXA2 might be necessary for NKX2-1 to promote NET. Indeed, simultaneous OE of NKX2-1 and FOXA2 in LNCaP cells drastically accelerated NET, with rapid induction of NE lineage markers, such as SYP, NCAM1, and BRN2, as early as D7 (**Fig.5D**). A majority of cells exhibited neurite extensions and “cluster” morphology at D14 of NKX2-1 and FOXA2 co-expressed cells (**Fig.S5C**). Furthermore, FOXA2 ChIP-seq revealed that FOXA2 rapidly completed its switch from luminal to NE enhancers as early as D14 in LNCaP+FOXA2+NKX2-1 cells with binding at NE enhancers even stronger than the D28 LNCaP+FOXA2 cells (**Fig.5E and S5D**). By contrast, most of FOXA2 remained at luminal enhancers in the D14 LNCaP+FOXA2 cells. Altogether, these data suggest that NKX2-1 facilitates and accelerates FOXA2-driven NET.

As NKX2-1 is selectively and highly expressed in NE tumors, we thought it may be an essential regulator of NEPC in general. To test this, we performed NKX2-1 KD in well-established NEPC cell line NCI-H660 and indeed observed an ablation of NE marker genes SYP and ASCL1 at both protein and mRNA levels (**Fig.5F**). Genetic alterations of TP53 and RB1 are found in 70% of clinical NEPC samples and their double knockout have been shown to drive NET [5, 6]. To investigate whether NKX2-1 is required for NEPC driven by these common genetic aberrations, we generated LNCaP cells with RB1 and TP53 double KD. NE markers, such as NCAM1, SYP, and SOX2, are up-regulated following the loss of RB1/P53, as expected (**Fig.5G**). Importantly, NKX2-1 is also induced in these cells, and concomitant depletion of NKX2-1 abolished NE marker gene expression, strongly supporting an essential function of NKX2-1 in mediating NET caused by genetic aberrations frequent in clinical samples. In addition, we analyzed 42D, an AR+/PSA- cell line that was derived *in vivo* upon resistance to AR antagonist enzalutamide, recapitulating clinical treatment-resistant PCa [20]. Interestingly, NKX2-1 OE remarkably increased the expression of NE markers such as SYP, ASCL1, and CHGB despite low FOXA2 expression in these cells, suggesting NKX2-1 as a general regulator of NET that likely also cooperates with other chromatin-pioneering factors (**Fig.5H**). Taken together, these data demonstrated that NKX2-1 is selectively up-regulated in NEPC and is an essential mediator of NET.

### FOXA2 induces regional DNA demethylation in NET models as well as patient-derived samples

We next attempted to understand the chromatin-initiating roles of FOXA2 in preparing the cells for NET. Previous studies of embryonic development have reported that FOXA2 is able to modulate regional DNA demethylation, a prerequisite for inheritable lineage-specific enhancer priming [56]. To test this in PCa, we first investigated global DNA methylation profiles in time-course LNCaP+FOXA2 cells and observed remarkable epigenetic differences between D2 and D28, including 16,470 hypermethylated regions (Hyper-DMRs) and 2,697 hypomethylated regions (Hypo-DMRs) at D28 (**Fig.6A)**. The DNA methylation pattern of D14, not surprisingly, was in between but more similar to D2, being consistent with the scMultiome data. Critically, FOXA2 binding at hyper-DMR regions gradually decreased over the time-course, whereas its binding at hypo-DMR regions increased, showing a negative correlation (**Fig.6B**). For example, there is drastic DNA hypomethylation at multiple CpG islands within the HOXB1-9 region in D28 cells, in concordance with increased FOXA2 binding, whereas the HOXB13 region was hypermethylated and showed less FOXA2 binding at D28 (**Fig.S6A**).

**Figure 6.**
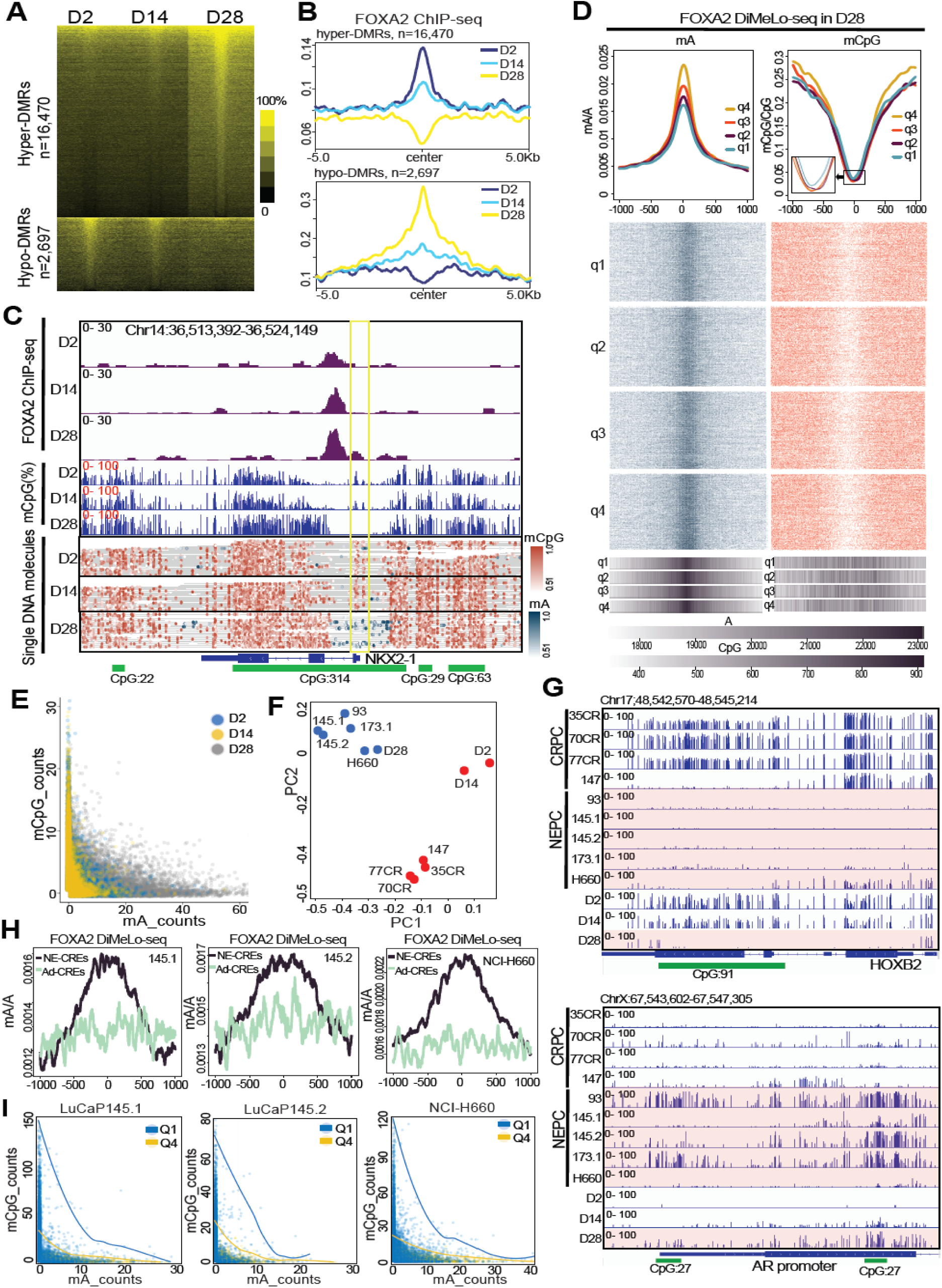
FOXA2 induces regional DNA demethylation in NET model as well as patient-derived samples. **A.** Heatmap showing the percentage of methylation around (+/-0.5kb) hyper- and hypo-DMRs in D2, D14, and D28 LNCaP+FOXA2 samples. **B.** Average intensity plots of FOXA2 ChIP-seq around hyper- and hypo-DMR sites shown in **A**. **C.** Integrative genomics viewer (IGV) view of FOXA2 ChIP-seq (**top**), mCpG (**middle**), and single DNA molecules of mCpG and mA (**bottom**) around the NKX2-1 gene in D2, D14, and D28 LNCaP+FOXA2 cells. Green box:CpG islands. Yellow box: NKX2-1 promoter. **D.** FOXA2 DiMeLo-seq was performed in D28 LNCaP+FOXA2 cells, mA and mCpG were called with a probability ≥ 0.5. Aggregate mA and mCpG curves (**top**) for each quartile were created with a 50-bp rolling window centered (±1kb) at D28-specific FOXA2 peaks, which were sorted into 4 quartiles with quartile 4 (q4) comprising the strongest peaks. Inset: a zoom-in version of the mCpG curves. Heatmap (**bottom**) shows mA and mCpG on single DNA molecules. A and CG base density (scale bar at the bottom) across the 2kb region of FOXA2 peaks within each quartile are shown in the 1D heatmaps. **E.** Correlation plots of mA and mCpG counts on the same DNA molecules in D2, D14, and D28 LNCaP+FOXA2 cells. **F.** PCA analyses of methylation profiles of CRPC/NEPC PDX and D2, D14, D28 LNCaP+FOXA2 samples using DMRs identified between D2 and D28 cells. **G.** IGV view of %mCpG in samples named on the left at HOXB2 (**top**) and AR promoters (**bottom**). CpG islands are shown in green. **H.** FOXA2 DiMeLo-seq showing FOXA2 binding (**mA/A)** at previously reported (Baca, Takeda et al. 2021) Ad-CREs and NE-CREs in NEPC tumors/cells. **I.** Correlation plots of mA and mCpG count on same DNA molecules in NEPC tumors/cells. Quartile 1 (Q1) comprises the weakest peaks, and quartile 4 (Q4) comprises the strongest peaks based on FOXA2 ChIP-seq signals in the respective samples. Blue and yellow lines indicate trend lines in Q1 and Q4, respectively.

To further examine whether FOXA2 binding induces regional DNA demethylation in PCa, we adapted the recently developed directed methylation with long-read sequencing (DiMeLo-seq) approach [57], which provides concurrent, base-pair level information on CpG methylation (mCpG) and TF binding on the same DNA molecules. Briefly, we first utilized FOXA2 antibody-tethered deoxyadenosine methyltransferase Hia5 to catalyze the formation of N6-methyl-deoxyadenosine (hereafter <u>mA</u>), which is rare in native human DNA, at FOXA2 binding sites and then used Oxford Nanopore Technology to directly sequence mA along with mCpG on the same DNA molecule. Interestingly, sharp peaks of DNA methylation were detected right at the NKX2-1 promoter in D0 cells and slightly decreased in D14, which were completely erased at D28 when abundant mA was detected around this region, indicating FOXA2 binding that is also supported by FOXA2 ChIP-seq data (**Fig.6C**). To analyze this globally, we stratified D28-specific FOXA2 binding sites into 4 quartiles based on FOXA2 ChIP-seq signal. As expected, FOXA2 peaks with stronger ChIP-seq signal (q4) had higher mA levels in D28 cells (**Fig.6D**). However, they were devoid of DNA methylation, indicated by slightly deeper and broader DNA methylation valleys. As a control, there were no mA and higher CpG methylation at these sites in D2 cells (**Fig.S6B**). Furthermore, at a single DNA molecule level, we observed a mutually exclusive pattern of mA and mCpG at each time point, with D28 cells containing DNA molecules with the highest mA but the lowest mCpG levels (**Fig.6E)**, supporting a dynamic FOXA2 binding and DNA demethylation process over the time-course.

To further validate FOXA2’s role in regulating regional DNA demethylation, we examined DNA methylation in an extensive list of prostate PDX tumors. We found that DNA methylation was significantly lower at Ad-CREs and NE-CREs, respectively, in CRPC and NEPC cells, being consistent with their respective luminal and NE enhancer activation (**Fig.S6C-E**). Moreover, PCA analyses using DMRs identified in our LNCaP+FOXA2 cells clearly separated CRPC from NEPC tumors. Importantly, we observed D2 and D14 samples being clustered together, adjacent to CRPC tumors, while D28 cells were closely grouped with NEPC samples, suggesting similar DNA methylation profiles between D28 cells and NEPC tumors (**Fig.6F**). For instance, HOXB2 gene, which was activated in D28 cells, was marked by dense CpG methylation in D2 cells and CRPC PDX tumors, which were completely erased in D28 cells as well as NEPC PDX tumors, whereas an opposite DNA methylation remodeling was observed at the promoter of luminal gene AR (**Fig.6G**). In addition, GO analyses showed that the genes near Hyper-DMRs were involved in epithelial cell function, whereas Hypo-DMRs genes are significantly enriched in embryo development and neurogenesis in our LNCaP+FOXA2 cells (**Fig.S6F**). Furthermore, PCA analyses using the top 200 each of Hypo- and Hyper-DMR genes grouped D2 LNCaP+FOXA2 cells adjacent to CRPC PDX and clinical CRPC samples, whereas D28 cells were closely clustered with NEPC samples (**Fig.S6G**). Next, we performed FOXA2 DiMeLo-seq for single-molecule analyses of FOXA2 binding and DNA methylation in NEPC cells wherein FOXA2 is expressed. Importantly, FOXA2 binding, reflected by the mA levels, was substantially more enriched at NE-CREs than Ad-CREs in all NEPC samples tested (**Fig.6H**). Analyses of mA and mCpG levels in single DNA molecules revealed a mutually exclusive pattern of FOXA2 binding and CpG methylation, as was confirmed in LNCaP+FOXA2 cells, strongly indicating that FOXA2 binding mediates regional DNA demethylation on the same DNA molecules (**Fig.6I**). Moreover, we found that regions of strong FOXA2 ChIP-seq signal (Q4) are associated with strong mA levels but diminished mCpG levels. In summary, our data showed that FOXA2 binding on the chromatin induces regional DNA demethylation, setting the stage for enhancer priming and lineage re-specification.

### NKX2-1 and FOXA2 recruit CBP/P300 to activate NE enhancers and NEPC tumor growth, which can be abolished by CBP/p300 inhibition

Now that we have demonstrated that FOXA2 OE induces DNA demethylation and NE enhancer priming and activation, we asked how the NE enhancers are activated. We analyzed FOXA2-interacting proteins in LuNE cells by mass spectrometry. Not surprisingly, among the top interactors were NKX2-1 and subunits of SWI/SNF that are recently reported to interact with FOXA1 in prostate and breast cancer cells [58, 59] (**Fig.7A**). Of high relevance, histone acetyltransferases CREBBP (CBP) and P300, the enzymes that catalyze H3K27ac, were also significantly enriched. Co-IP with FOXA2 or NKX2-1 antibodies confirmed their interaction with both CBP and P300 in LuNE cells (**Fig.7B**). This endogenous protein complex was further validated in the well-established NEPC cell line NCI-H660 (**Fig.7C**). To investigate the function of this complex, we focused on FOXA2 binding sites. Time-course FOXA2 ChIP-seq data revealed 5 clusters, 3 of which (C2, C4 and C5) are NE enhancers, showing increased binding upon NET (**Fig.7D, left**). Of note, NKX2-1 showed enriched binding at NE enhancers, and this binding is specific to D21 and D28 cells, concurrent with major FOXA2 cistrome switch from luminal to NE enhancers. ChIP-seq in LuNE cells demonstrated enriched P300 and H3K27ac at NE enhancers, which were abolished by either FOXA2 or NKX2-1 KD (**Fig.7D, middle**), suggesting that both FOXA2 and NKX2-1 are required for P300 recruitment to these enhancer elements to catalyze H3K27ac. Moreover, H3K27ac at NE enhancers was lost upon P300 inhibition, either by p300 KD or using pharmacological inhibitors, such as CCS1477 (**Fig.7D, right**). These data support that FOXA2 and NKX2-1 cooperate to active NE enhancers through P300.

**Figure 7.**
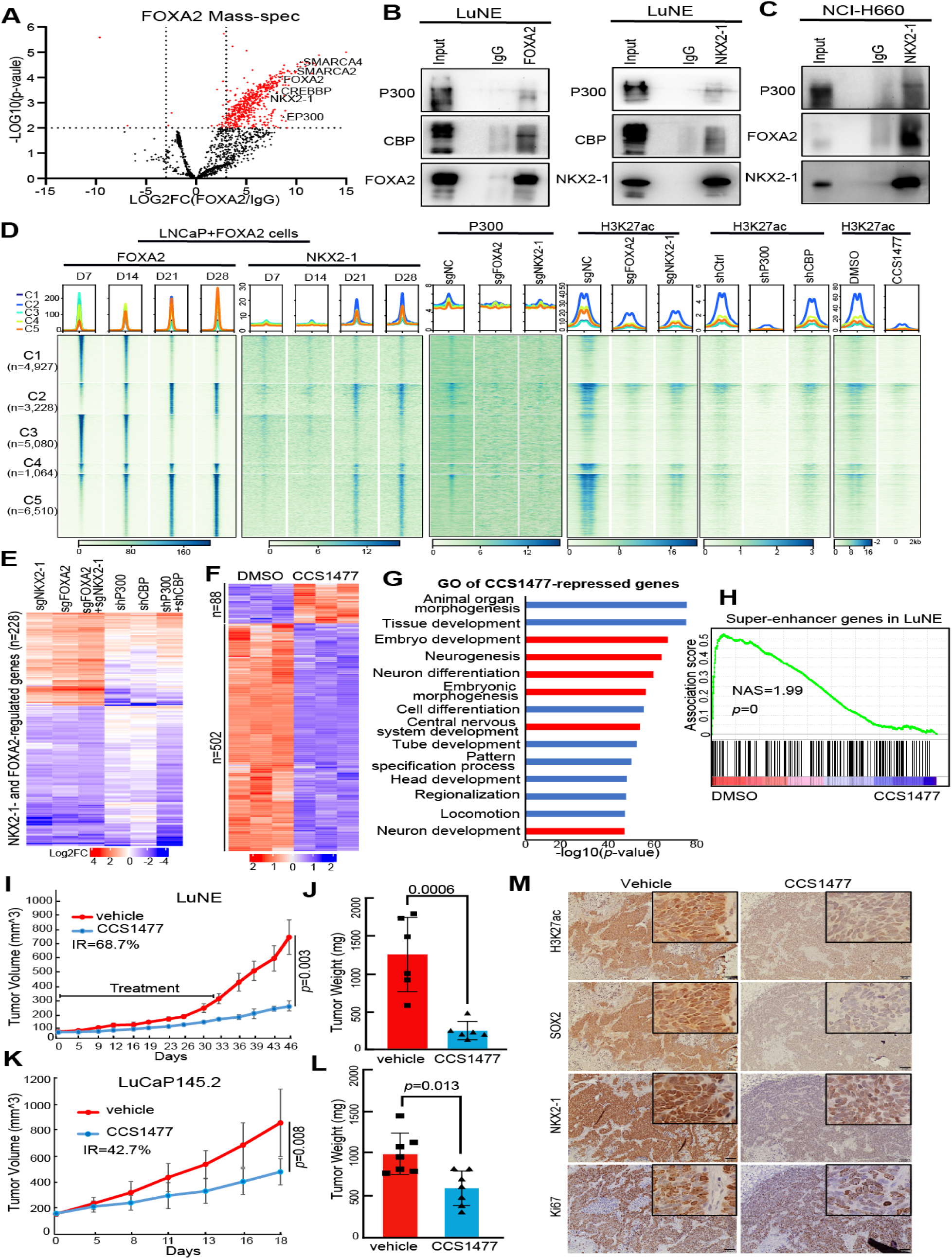
NKX2-1 and FOXA2 recruit CBP/P300 to activate NE enhancers and tumor growth, which can be abolished by CBP/p300 inhibition. **A.** Volcano plot showing FOXA2-interacting proteins in LuNE cells by mass-spec (pooled data from 3 Co-IP replicates). The x-axis represents log2(FC), and the y-axis represents −log10 of *p*-values by *t*-test. Each dot represents a protein, with red dots: log2FC ≥3 and *p* < 0.01. **B.** Co-IP showing that FOXA2 and NKX2-1 interact with CBP and P300 in LuNE cells. **C.** Co-IP showing that NKX2-1 interacts with FOXA2 and P300 in NCI-H660 cells. **D.** Heatmaps showing indicated ChIP-seq intensity centered (±2kb) around the 5 clusters of FOXA2-binding sites identified in the time-course LNCaP+FOXA2 cells. Scale bar: enrichment intensity. **E.** Heatmap showing that genes induced by FOXA2 and NKX2-1 (FC>2 and adjusted *p*<0.05) are regulated by P300/CBP in LuNE cells. Data shown is the log2FC of sgNKX2-1, sgFOXA2, or sgFOXA2+sgNKX2-1 relative to sgNC, and of shP300, shCBP, or shP300+shCBP to shCtrl. **F.** LuNE cells were treated with DMSO or CCS1477 (250nM) for 72h. Differentially expressed genes were identified by DESeq2 with FC≥2, adjusted *p*<0.05. Color bar: z-scores. **G.** GO analysis of CCS1477-repressed genes. Top enriched molecular concepts are shown on Y-axis, while X-axis indicates enrichment *P* values calculated by hypergeometric test. **H.** GSEA of SE-associated genes in LuNE cells treated with DMSO or CCS1477. **I-J**. Tumor growth curve (**I**) and weight (**J**) of LuNE xenograft tumors treated with vehicle or CCS1477. Data are mean±SD, and *p* values by *t* test. **K-L**. Tumor growth curve (**K**) and weight (**L**) of LuCaP145.2 xenograft tumors treated with vehicle or CCS1477. Data are mean±SD, and *p* values by *t* test. **M**. Representative IHC images of H3K27ac, SOX2, NKX2-1, and Ki67 in LuCaP145.2 xenograft tumors treated with vehicle or CCS1477. Scale bar, 30 µm. Insets are shown at higher magnification.

To determine whether NKX2-1 and FOXA2-driven neuronal transcription programs are dependent on P300/CBP in LuNE cells, we performed RNA-seq in LuNE cells with KD of control, P300, CBP, or both. Importantly, we found that most NKX2-1/FOXA2-regulated genes were also regulated by CBP or P300 KD in a similar fashion, suggesting a strong dependence of this NE gene expression pattern on the complex (**Fig.7E and Fig.S7A**). Moreover, GO analysis of P300/CBP-induced genes in LuNE cells illustrated significant enrichment in tissue development, neurogenesis, embryo development, and cell proliferation, similar to FOXA2/NKX2-1-induced genes (**Fig.S7B**). Concordantly, KD of either NKX2-1, FOXA2, P300 and, to a lesser extent, CBP all significantly inhibited LuNE cell colony formation (**Fig.S7C**), revealing a vulnerability of NEPC cells to therapeutics targeting this pathway.

We thus decided to evaluate the therapeutic potential of CCS1477, an orally active and selective inhibitor of P300/CBP bromodomain that is currently in clinical trials in CRPC patients via targeting AR [60]. To test whether this drug is equally effective in AR-NEPC models, we treated LuNE cells with CCS1477 and noticed remarkable suppression of gene expression (**Fig.7F**). GO analysis of CCS1477-repressed genes identified neurogenesis and embryonic development processes similar to CBP/P300 KD, indicating lineage-specific therapeutic targets (**Fig.7G**). To further affirm that these effects are due to CCS1477-targeting NE enhancers, we performed super-enhancers analyses based on H3K27ac ChIP-seq in LuNE cells using ROSE [61]. We identified 217 super-enhancers, and their associated genes indeed include developmental regulators, such as HOXA5/9, ZIC2/5, EZH2, and cell cycle regulators including E2F3 (**Fig.S7D**). Gene Set Enrichment Analysis (GSEA) validated that these super-enhancer-associated genes are strikingly enriched for suppression by CCS1477 (**Fig.7H**).

Next, we sought to investigate how well CCS1477 is able to control NEPC growth. Treating LuNE cells with increasing doses of CCS1477, we found it massively inhibited colony formation at only 0.1μM concentration and depleted LuNE cell growth at 0.5μM (**Fig.S7E**). Similarly, CCS1477 achieved a strong killing effect at a dose lower than 0.5μM in two additional NEPC models, LuCaP145.2 and NCI-H660 (**Fig.S7F**). As controls, we treated benign prostatic hyperplasia cell line BPH-1 and normal prostate epithelial cell line RWPE-1 with increasing doses of CCS1477 up to 500nM and did not observe apparent, precluding un-targeted toxicity of this drug (**Fig.S7G**). To examine the efficacy of CCS1477 on NEPC tumor growth *in vivo*, we performed LuNE xenograft in SCID mice. Once tumor size reached 70mm^3, mice were randomized to receive treatment with vehicle (5% DMSO:95% methylcellulose) or CCS1477 (20 mg/Kg) daily by oral gavage. We started to see apparent growth reduction after 3 weeks of treatment. Interestingly, despite treatment ending at day 33, tumor growth continued to be controlled by CCS1477, with an inhibition rate (IR) of 68.7% (*p*=0.003) at day 46 when the experiment was terminated (**Fig.7I**). There was a remarkable difference in endpoint tumor weight (*p*=0.0006) and tumor sizes (**Fig.7J and Fig.S7I**), while the body weight of the mice was not significantly affected by the drug (**Fig.S7H**). Similar tumor-inhibitory effects of CCS1477 were also observed in NEPC PDX (LuCaP145.2) tumors (**Fig.7K-L and Fig.S7J**). IHC analyses of endpoint tumors confirmed that CCS1477 treatment drastically reduced H3K27ac, Ki67, and NE transcription factors NKX2-1 and SOX2 (**Fig.7M and Fig.S7K**). In summary, these data support that CBP/P300 plays important roles in NE enhancer activation, which can be effectively targeted to mitigate NEPC growth *in vitro* and *in vivo* using CCS1477.

## DISCUSSION

The 3D chromatin architecture as an instructive force of transcriptional regulation has been demonstrated to be cell type-specific and critical for cell identity determination [28-30, 62]. CRPC and NEPC are two different subtypes of advanced PCa, which have distinct epigenetic and transcriptional programs. Consistent with this, our data showed the 3D genome conformation was remarkably reorganized in NEPC compared to CRPC tumors, despite substantial heterogeneity within the group. This chromatin reorganization leads to the loss of luminal E-P loops and the formation of new E-P loops and TADs that favor an NE program. We demonstrated that this observation in patient-derived samples can be recapitulated in isogenic cells undergoing NET *in vitro* upon FOXA2 OE. We found new E-P loops mediated by enhancer-bound FOXA2 and promoter-bound NKX2-1, emerging at D14 and drastically increasing at D21 and D28. The initial E-P loop formation may be mediated by cofactor oligomerization between FOXA2 and NKX2-1, which may be further strengthened at later time points as the NE enhancers get activated by CBP/P300, attributing to condensate formation. Previous evidence suggests that master transcription factors, as anchor proteins, are able to mediate long-range DNA interactions through mechanisms, including cofactor oligomerization and condensate formation [30, 62]. Moreover, CBP/P300 proteins are characterized by extensive intrinsically disordered regions, which are known to promote multivalent interactions and the formation of subnuclear condensates [63, 64]. Although our study focused on demonstrating this mechanism using NKX2-1 and FOXA2 model, we believe 3D chromatin rewiring is a general mechanism that underlines NET and permanently stabilizes the NEPC lineage, as a 3D chromatin architecture favoring NE program was observed in NEPC tumors with or without FOXA2 expression (**Fig.S1A**).

The expression and functions of neural TF NKX2-1 have not been well characterized in PCa. In this study, we report that NKX2-1 is selectively and highly expressed in AR-/NE+ NEPC tumors. We demonstrated that NKX2-1 is a direct transcriptional target of FOXA2 and is indispensable for FOXA2-driven NET. Mechanistically, FOXA2, as a pioneer factor, binds to DNA to mediate regional DNA demethylation and initiate local chromatin opening, leading to NE enhancer priming at early time points. Once NKX2-1 is induced by FOXA2, promoter-bound NKX2-1 interacts with enhancer-bound FOXA2 by forming E-P loops, further increasing FOXA2 recruitment to NE enhancers and leading to NE enhancer activation and 3D genome rewiring. A recent study has shown that FOXA2 is recruited by organ-specific TFs to induce enhancer priming and enforce lineage-specific gene expression during pancreatic, hepatic, and lung development [65]. The recruitment of lineage-specific TFs can, in turn, enhance FOXA protein binding, especially at unprimed enhancers, to enforce organ cell type-specific gene expression [65], which is consistent with our observation that NKX2-1 facilitates FOXA2 binding at NE enhancers for ultimate activation in NEPC. Although our current study deciphers the in-depth mechanism using the FOXA2-driven NET model, the mechanism identified here may be generable, as FOXA2 and NKX2-1 are co-expressed in most NEPC patients. Whereas there is a small number of NEPC that only expresses NKX2-1, it might function through other chromatin pioneering factors, such as ASCL1. Indeed, we further validated that NKX2-1 is essential for NET across various models, strongly supporting NKX2-1 as a robust promoter of NEPC progression.

The latest advance of DiMeLo-seq using long-read sequencing provides a powerful tool to study TF binding and DNA methylation on the same DNA sequencing reads [57]. We found that FOXA2 binding leads to regional DNA demethylation on the same DNA molecules. Our finding in NEPC is consistent with recent reports that FOXA2 interacts with TET1 dioxygenase in endodermal lineage intermediates and mediates regional DNA demethylation (spanning ±800bp around the FOXA2-binding sites) that is critical for liver and pancreatic endocrine cell specification [56, 66]. In addition, in prostate cancer cells, we have previously demonstrated an interaction between FOXA1 and TET1 protein to mediate DNA demethylation [67]. Considering the sequence and functional similarity between FOXA1 and FOXA2, it may be assumed that TET1 is also involved in FOXA2-induced DNA demethylation reported in this study. Studies of embryonic development have suggested that FOXA2, as a chromatin-pioneering TF, may prime lineage-specific enhancers by mediating regional DNA demethylation and depositing H3K4me1 *de novo*, while subsequent recruitment of lineage-inductive TFs leads to enhancer activation [56, 68-73]. Our study demonstrated the important role of FOXA2 in initiating an epigenetic memory of NE cellular identity and lineage specification. While how DNA demethylation leads to enhancer priming and activation is beyond the scope of the current study, FOXA1 is known to recruit MLL3 to induce H3K4me1 at target enhancers [70].

Presently, there is no effective treatment for NEPC except platinum-based chemotherapy. Therefore, new therapeutic approaches are urgently needed. Our studies found that FOXA2 and NKX2-1 turn on NE lineage transcription programs by recruiting CBP/P300 to NE enhancers. Several CBP/P300 inhibitors have been developed, including A485 and CCS1477, with the latter already in clinical trials for late-stage, metastatic CRPC due to its ability to target AR and c-Myc [60]. We demonstrated that, in NEPC cells, CCS1477 significantly inhibited NEPC tumor growth by suppressing NE-specific super-enhancers and gene expression. Thus, CCS1477 may be beneficial in inhibiting both CRPC and NEPC by targeting distinct luminal and NE enhancers that are respectively important for their growth. CCS1477 may be useful in preventing the development of treatment-resistant NEPC tumors in CRPC patients or inhibiting NEPC tumor growth when used later in the disease time course. It will be of great interest to test the efficacy of CCS1477 in CRPC and NEPC patient populations in future clinical trials and compare the downstream pathways affected.

## MATERIALS AND METHODS

### Ethics statement

Our research complies with all relevant ethical regulations. Mouse handling and experimental procedures were approved by the Institutional Animal Care and Use Committee at Northwestern University in accordance with the US National Institutes of Health Guidelines for the Care and Use of Laboratory Animals and the Animal Welfare Act.

### Cell lines, chemical reagents, and antibodies

Prostate cancer cell lines LNCaP, 22Rv1, DU145, NCI-H660, human benign prostatic hyperplasia cell line BPH-1, human normal prostate epithelial cell line RWPE-1and human embryonic kidney cell line HEK293T cells were obtained from American Type Culture Collection (ATCC). LNCaP, 22Rv1, DU145, BPH-1, and HEK293T were maintained in either RPMI1640 or Dulbecco’s modified Eagle’s medium (DMEM) with 10% fetal bovine serum (FBS), 1% penicillin and streptomycin. RWPE-1 cells were maintained in Keratinocyte Serum-Free Medium (K-SFM) with 0.05 mg/ml BPE and 5 ng/ml EGF. NCI-H660 cells were maintained in ATCC-formulated RPMI-1640 Medium (Catalog No.30-2001) with 0.005 mg/ml Insulin, 0.01 mg/ml Transferrin, 30nM Sodium selenite, 10 nM Hydrocortisone, 10 nM beta-estradiol, 5% fetal bovine serum. 42D cells were maintained in RPMI1640 with 5% fetal bovine serum (FBS), 1% penicillin and streptomycin, and 10μM ENZ for all experiments. All the cells were authenticated within 6 months of growth, and cells under culture were frequently tested for potential mycoplasma contamination. CCS1477 (Catalog No. CT-CCS1477) was purchased from Chemietek. All antibodies used in this study are listed in **Supplementary Table 2.**

### Constructs and Lentivirus Infection

Human FOXA2 coding region (CDS) was first amplified using cDNA from PC-3 cells as a template, NKX2-1 and mouse Foxa2 CDS were amplified using NKX2-1 (Addgene, #119173) and mFoxa2 (Addgene, #33014) as template respectively and then cloned into plenti CMV Neo DEST (705-1) (Addgene, #17392) by In-Fusion HD Cloning Kits (TaKaRa, Cat#638948). FOXA2, NKX2-1, RB1 and TP53 gRNAs were cloned into lentiCRISPR v2 (Addgene, #52961). The shRNAs targeting CBP and P300 were cloned into pLKO.1-TRC lentiviral vector (Addgene, #10878). The primers and oligonucleotides used in this study are listed in **Supplementary Table 3**, and all the plasmids were verified by Sanger sequencing. For the generation of lentivirus, HEK293T cells were transfected with psPAX2 and pMD2G with the target gene at a ratio of 2:1:1. The supernatant containing lentiviruses was harvested at 48 h after transfection and filtered through a 0.45μm filter. Lentiviruses, supplemented with 8μg/mL polybrene, were used to infect human PCa cells. For most knockdown experiments with shRNAs or gRNAs, cells were harvested on day 5 after infection. For organoid infection, the single cells were mixed with lentivirus-containing polybrene with a final concentration of 8μg/mL and then centrifuged for 1 h at 600g at room temperature. Cells were subsequently placed at 37°C, 5% CO2, for 6 h to recover before being plated in Matrigel.

### Co-immunoprecipitation (Co-IP)

Nuclear fraction was used for all Co-IP experiments in this study. Nuclear proteins were isolated as previously with some modifications [46, 74]. Briefly, cells were resuspended in buffer A (10 mM HEPES, pH 7.9, 10 mM KCl, 1.5 mM MgCl2, 0.34 M sucrose, 10% glycerol, 1 mM dithiothreitol, 1mM EDTA, 1 × Roche protease inhibitor cocktail) and incubated on ice for 10 min. Then, the TritonX-100 with a final concentration at 0.1% was added to the cell suspension to extract cytoplasmic fraction. After washing once with buffer A, the nuclei were incubated in buffer B (10mM HEPES, pH7.9, 10% glycerol, 1 mM EDTA, 1.5 mM MgCl2, 300 mM NaCl, 0.5% NP 40, 1 mM dithiothreitol, 1 × Roche protease inhibitor cocktail) for 30 min on ice to isolate the nuclear fraction. The nuclear fraction was first pre-cleared with protein G or A-magnetic beads at 4 °C for 2 h, followed by incubation with the corresponding antibody overnight. Dynabeads protein G (Life Technologies) was added the next day and incubated for 1 h at 4 °C. The beads/protein complex was washed four times with IP wash buffer (10mM HEPES, pH7.9, 1 mM EDTA, 1.5 mM MgCl2, 150 mM NaCl, 0.5% NP 40, 1 mM dithiothreitol, 1 × Roche protease inhibitor cocktail) and eluted with 30 µl of 2 × SDS sample buffer and subjected to western blot (WB) analysis using the corresponding antibody. All antibodies used in this study are listed in **Supplementary Table 2**.

### 3D organoid cell culture

LuCaP 145.2 PDX-derived organoids were generated as previously described [75]. Briefly, LuCaP145.2 tumors were minced into small pieces (∼1 mm3) and digested in 5 mg/ml collagenase type II (Gibco, Cat#17101-015) with 10 μM Y-27632 dihydrochloride (Tocris, Cat#1254) in a 15-ml Falcon tube for 1–1.5 h at 37 °C. Digested tissues were washed once with Advanced DMEM/F12 medium containing penicillin/streptomycin, 10 mM HEPES and 2 mM GlutaMAX (adDMEM/F12 +/+/+), then resuspended in 5 mL of TrypLE Express (Gibco, 12604-021), and further digested at 37 °C for 15 min. After digestion, cells were washed once with adDMEM/F12 +/+/+ and counted with a hemocytometer. A total of 20,000 cells were mixed with 50% Matrigel and plated in a 12-well tissue culture plate, and placed in a CO2 incubator (5% CO2, 37 °C) for 15 min to allow Matrigel to solidify. Prewarmed human complete organoids medium plus 10 μM Y-27632 dihydrochloride was gently added to cells, and cells were maintained in a CO2 incubator (5% CO2, 37 °C). The medium was changed every 3 days. The organoids were passaged biweekly by 1:3.

### Colony formation assay

For colony formation assay, LuNE cells with indicated gene alteration (5 ×10^3^ cells per well), RWPE-1, and BPH-1 (2 ×10^3^ cells per well) cells were seeded in 12-well plates. The cells were fixed by 4% paraformaldehyde after 2 weeks of growth and stained with 0.05% crystal violet. The colonies were imaged with ChemiDoc (BIO-RAD).

### Mass spectrometry analysis

Chromatin fraction was used for the mass spectrometry experiments in this study. Chromatin proteins were isolated as previously with some modifications [46, 74]. Briefly, cells were resuspended in buffer A (10 mM HEPES, pH 7.9, 10 mM KCl, 1.5 mM MgCl2, 0.34 M sucrose, 10% glycerol, 1 mM dithiothreitol, 1 × Roche protease inhibitor cocktail) and incubated on ice for 10 min. Then, the final concentration of 0.1% TritonX-100 was added to the cell suspension and vortexed for 15 s, and spun down at 4 °C for 5 min at 1,000g. The supernatant was kept as a cytoplasmic fraction. The nuclei pellet was washed once with buffer A, and then resuspended in buffer C (3 mM EDTA, 75 mM NaCl, 0.1% TritonX-100, 1 mM dithiothreitol, protease cocktail) for 30 min on ice to remove chromatin free proteins. Insoluble chromatin was resuspended in buffer D (10mM HEPES, pH7.9, 10% glycerol, 1 mM EDTA, 1.5 mM MgCl2, 300 mM NaCl, 0.5% NP 40, 1 mM dithiothreitol, 0.5U/μl TurboNuclease (Accelagen, Cat#N0103M), 1 × Roche protease inhibitor cocktail) and incubated at 4 °C for 30 min in a rotor with 300 r.p.m. The chromatin was spun down for 10 min at 12,000g at 4 °C, and the supernatant was saved as the chromatin fraction. The chromatin fraction was first pre-cleared with protein A-magnetic beads at 4 °C for 2 h, followed by incubation with FOXA2 (Abcam, Cat#ab108422) or IgG (Sigma, cat#12-370) antibody overnight. Dynabeads Protein A (Life Technologies) were added the next day and incubated for 1 h at 4 °C. The beads/protein complex was washed four times with IP wash buffer (10mM HEPES, pH7.9, 1 mM EDTA, 1.5 mM MgCl2, 150 mM NaCl, 0.5% NP 40, 1 mM dithiothreitol, 1 × Roche protease inhibitor cocktail). The beads were eluted with 1xSDS sample buffer and subjected to SDS–PAGE. Protein bands were excised and subjected to mass spectrometry analysis using the Orbitrap Velos Pro system. The SEQUEST was used for protein identification and peptide sequencing. The data in Fig.7A was analyzed as previously [76]. Briefly, the sum of all peptide intensities of one protein in triplicate was log2 transformed, normalized by average, then missing peptide intensities were imputed from a Gaussian distribution of width 0.3 centered at the sample distribution mean minus 2.5x the sample standard deviation. The criterion for statistically significant differential interactions was calculated by unpaired two-side *t*-test with a *p*-value less than 0.01 and fold change (FOXA2 vs. IgG) great than 8.

### RNA extraction, RT–qPCR, and RNA-seq

RNA was extracted using the nucleospin RNA kit (Takara) according to the manufacturer’s recommended protocol. Then, 500 ng of RNA was reverse transcribed into complementary DNA (cDNA) using the ReverTra Ace qPCR RT Master Mix kit (Diagnocine) according to the manufacturer’s recommended protocol. qPCR was performed with 2X Universal SYBR Green Fast qPCR Mix (Abclonal, Cat#RK21203) and QuantStudio 3 (Thermo Fisher). All primers used here are listed in **Supplementary Table 3**. For RNA-seq, total RNA was isolated as described above and performed in triplicate. RNA-seq libraries were prepared from 0.5 μg of high-quality DNA-free RNA using NEBNext Ultra RNA Library Prep Kit, according to the manufacturer’s instructions. The libraries passing quality control (equal size distribution between 250 and 400 bp, no adapter contamination peaks, no degradation peaks) were quantified using the Library Quantification Kit from Illumina (Kapa Biosystems, KK4603). Libraries were pooled to a final concentration of 10 nM and sequenced pair-end using the Illumina Novaseq 6000.

### ChIP, ChIP–seq, and ATAC–seq

ChIP and ChIP–seq were performed using the previously described protocol with the following modifications [46, 74]. For FOXA2, NKX2-1, H3K27ac, and H3K4me1 ChIP, PCa cells were crosslinked with 1% formaldehyde (Thermo Fisher, Cat#28908)) for 10 min at room temperature with gentle rotation and then quenched for 5 min with 0.125 M glycine. A total of 10 million cells were used for each FOXA2, NKX2-1 ChIP, and 5 million cells were used for each H3K4me1 or H3K27ac ChIP. Chromatin was sonicated to an average length of 200–600 bp using an E220 focused ultrasonicator (Covaris). Supernatants containing chromatin fragments were pre-cleared with protein A agarose beads (Millipore) for 40 min and incubated with a specific antibody overnight at 4 °C on a rotator (antibody information is listed in **Supplementary Table 2**). Then we added 50 μl of protein A agarose beads and incubated for 2 h. Beads were washed twice with 1 × dialysis buffer (2 mM EDTA, 50 mM Tris-Cl, pH 8.0) and four times with ChIP wash buffer (100 mM Tris-Cl, pH 9.0, 500 mM LiCl, 1% NP40, 1% deoxycholate). The antibody/protein/DNA complexes were eluted with elution buffer (50 mM NaHCO3, 1% SDS), the crosslinks were reversed, and DNA was purified with DNA Clean & Concentrator-5 kit (ZYMO Research). For H3K27ac ChIP in P300, CBP, NKX2-1, or FOXA2 KD and CCS1477-treated cells, we used Drosophila S2 spike-in normalization method and followed the protocol from Active Motif. ChIP–seq libraries were prepared from 3–5 ng of ChIPed DNA using NEBNext Ultra II DNA Library Prep Kit (NEB, E7645S), according to the manufacturer’s instructions. Post-amplification libraries were size selected between 250 and 450 bp using Agencourt AMPure XP beads from Beckman Coulter and were quantified using the Library Quantification Kit from Illumina (Kapa Biosystems, KK4603). Libraries were pooled to a final concentration of 10 nM and sequenced single end using the Illumina HiSeq 4000 or Novaseq 6000.

For FOXA2, NKX2-1, H3K4me1, H3K4me3 and H3K27ac ChIP in LuCaP PDX, double crosslinking was performed. 50 mg of homogenized LuCaP PDX tumor tissues were crosslinked with 2 mM Disuccinimidyl glutarate (DSG, Pierce) for 10 min at room temperature, followed by 1% formaldehyde for 10 min. Crosslinked cells were then quenched with 0.125 M glycine for 5 min at room temperature. Chromatin shearing, immunoprecipitation, and library preparation were performed as described above.

The assay for transposase-accessible chromatin using sequencing (ATAC–seq) was performed as previously reported [77]. Briefly, 50,000 cells were washed once with 1 × PBS and resuspended in 100 μl of lysis buffer (10mM Tris-HCl, pH 7.4, 10mM NaCl, 3mM MgCl2, 0.1% NP 40, 0.1% Tween-20, 0.01% Digitonin) for 3min on ice, followed by adding 1ml of wash buffer (10mM Tris-HCl, pH 7.4, 10mM NaCl, 3mM MgCl2, 0.1% Tween-20) to remove cytoplasm and mitochondria fraction. The nuclei pellet was resuspended in 25 μl transposition mix (12.5 μl of 2x TD buffer, 2.5 μl of Tn5 Transposome, 8.25 μl of 1xPBS, 0.25 μl of 1% digitonin, 0.25 μl of 10% Tween-20, 1.25 μl H2O) and incubated at 37 °C for 30 min in a thermomixer with 300 r.p.m. mixing. The tagmented DNA was purified using the DNA Clean & Concentrator-5 kit (ZYMO Research). Libraries were amplified, and adapter dimers and primer dimers were cleaned up. Paired-end reads (50 bp) were sequenced using an Illumina HiSeq 4000. **Bulk RNA-seq analysis**

Human prostate cancer cell RNA-seq reads were mapped to NCBI human genome GRCh38. Raw counts of genes were calculated by STAR. FPKM values (fragments per kilobase of transcript per million mapped reads) were calculated by in-house Perl script. Differential gene expression analysis was performed by the DESeq2 (1.26.0), which uses shrinkage estimation for dispersions and fold changes to improve the stability and interpretability of estimates.

Differentially expressed genes across the time course were performed using the likelihood ratio test option, otherwise, the standard pairwise comparisons were used. The AR signature and NE signature genes used in Fig.2B were from a previous study [3]. The scatterplot was generated using GSVA [52]. RNA-seq data of prostate cancer patients was downloaded from GSE126078[45].

### ChIP–seq analysis

ChIP–seq reads were aligned to the Human Reference Genome (assembly hg19) using Bowtie2 2.0.5. FastQC/0.12 was used to check quality control. The adapter was trimmed by Trim Galore/0.6.5. Non-uniquely mapping and redundant reads, and blacklists were removed. The remaining reads were used to generate binding peaks with MACS2 with a q-value (FDR) threshold of 0.01. For replicates, the peaks were merged for downstream analysis. Weighted Venn diagrams were created by the R package Vennerable. Heatmap views of ChIP-seq were generated by deepTools. Motif analyses were performed using HOMER. Genomic distribution of ChIP-seq binding sites was generated by the R Bioconductor package ChIPseeker/1.36. For FOXA2 clusters in Fig.7D, the sample size was down to the same size, peaks of FOXA2 with signal value greater than 10 were retained for generating heatmap. For H3K27ac ChIP-seq analysis in Fig.7D, we used Drosophila S2 spike-in normalization strategy and followed the protocol from Active Motif.

### ATAC-seq analysis

The raw data were processed with the ENCODE ATAC-seq pipeline (https://github.com/ENCODE-DCC/atac-seq-pipeline). In short, the reads were trimmed, filtered, and aligned against hg38 using Bowtie2. PCR duplicates and reads mapped to the mitochondrial chromosome or repeated regions were removed. To correct for the Tn5 transposase insertion, mapped reads were shifted +4/-5. Peak calling was performed using MACS2, with *a p*-value < 0.01 as the cutoff. Reproducible peaks from two biological replicates were defined as peaks with Irreproducibility Discovery Rates (IDR) < 0.05. A list of consensus peaks was created from non-overlap peaks based on summits extended by 250bp. Reads falling into consensus peaks were quantified by RSubread. For Fig.3A, differential ATAC-seq peaks were identified by DESeq2 using the LRT test. Peaks were filtered with padj < 0.001 and clustered using ComplexHeatmap.

### Single-cell sequencing sample preparation and analysis

A total of 5,000 of D0, D14, and D21 LNCaP+FOXA2 cells were used for single-cell Multiome ATAC + Gene Expression assay, the multiome libraries were prepared using 10X genomics Chromium Next GEM Single Cell Multiome ATAC + Gene Expression Reagent Kits (Cat# 1000285) per manufacturer’s protocol. All the single-cell libraries were sequenced in a NovaSeq 6000 with an average depth of 30,000 reads per cell. Raw sequencing data were preprocessed and aligned to hg38 using Cell Ranger ARC (2.0.2). Low-quality cells with low UMI counts and high mitochondrial ratios were filtered out, and the sequence metrics were updated in **Supplementary Table 4.** Each time point was analyzed individually at the expression and accessibility modality, then combined across time points within each modality using Seurat (4.3.0) and Signac (1.6.0) in R (4.0.3). For each time point, we performed quality control on ATAC fragments, RNA counts, nucleosome signal, and transcription start site enrichment metrics resulting in 2110, 1739, and 2344 cells for D2, D14, and D28, respectively. The RNA modality was normalized using SCTransform. ATAC peaks were called using MACS2 and quantified. Counts were normalized using latent semantic indexing. These were performed for each condition separately.

To correct for batch effects, integration of the expression modality was done using the Seurat V3 integration pipeline, using reciprocal PCA to identify anchors with 3000 integration features. UMAP was constructed using the first 30 PCs. To visualize the gene expression, we used LogNormalized data. AR and NE signatures were calculated using ‘AddModuleScore’. AR+ was defined as ≥ 0.6, which is the mean of AR normalized expression in D21. ARS+ and NES+ cells are defined as cells with a signature score ≥ 0. Monocle3 (0.2.3.0) was used to learn the trajectory graph and calculate the pseudotime. Velocyto (0.17.17) was used to generate spliced and unspliced matrices. RNA velocities were projected into the UMAP produced from earlier steps in Seurat, using Pyplot (matplotlib 3.7.0). Integration of the chromatin accessibility was done by first finding a common peak set between all the conditions, counting features, and re-normalizing. Integration anchors were found using “rlsi” using 2:50 dimensions. AR and NE signatures for the chromatin modality were performed by first finding peaks that fall within 5kb upstream and in the gene body of the respective genes. The scores were obtained using Signac’s ‘AddChromatinModule’, which calculates chromVAR deviations.

Patient samples’ scRNA-seq from Cheng et al. [53] was downloaded as bam files, converted into fastq files, and re-run using CellRanger (6.1.2). CellRanger aggr was run on the samples to recapitulate the original paper’s pipeline. We assigned cell types based on the markers mentioned in the paper. Our LNCaP+FOXA2 time course and the patient samples’ scRNA-seq data were normalized using SCTransform. Integration was performed using the SCTransform integration pipeline. After integration with the entire population, the non-epithelial cells and basal cells were removed, and a new UMAP was generated. The pseudotime was calculated with a node in the primary PCa population chosen as the starting cell.

### Hi-C analysis

FOXA2-D0, D14, D21, and D28 LNCaP cells, NCI-H660 cells were fixed with 2% formaldehyde (final concentration) at room temperature for 10 min, whereas tumor tissues from LuCaP PDX were fixed with 2% formaldehyde at room temperature for 20 min, followed by adding 2.5 M glycine to a final concentration of 0.2 M to quench the reaction. Hi-C libraries were generated using the Arima-HiC+ kit (Arima Genomics, A510008) per manufacturer’s protocol. The Hi-C libraries were sequenced with paired-end 2 × 150 bp using Illumina NovaSeq6000 or NovaSeq X Plus. The sequence metrics were updated in **Supplementary Table 1.** Hi-C data were processed into .mcool maps using the runHiC (0.8.6) pipeline, with chromap as the aligner. A/B compartments were called using cooltools (0.5.4) eigs_cis function with GC content phasing at 100kb resolution. TADs were called using cooltools insulation scores function. Loops were called using mustache (1.2.0) at 10kb resolution using default parameters. Each NEPC was compared to each CRPC using diffMustache with 0.001 cutoff. Each sample specific loops were defined as loops present in at least 2 of the comparisons. NEPC-or CRPC-specific loops are present in at least 2 NEPC or CRPC samples respectively. APA plots were made with coolpup.py (1.0.0). FOXA2 and NKX2-1 anchored loops were annotated based on the presence of matched ChIP-seq binding sites within the anchors.

### Reduced representation methylation sequencing (RRMS) using Nanopore Technology

The RRMS libraries were performed using the protocol from Oxford Nanopore. Briefly, the genomic DNA of LuCaP PDX was extracted with Quick-DNA Miniprep Plus Kit (Zymo, D4068) and fragmented to average size at 8 kb with g-TUBE™ (Covaris, 520079). The DNA libraries were prepared using Ligation Sequencing Kit (ON SQK-LSK110) per the manufacturer’s protocol. Sequencing was performed on an Oxford Nanopore GridiON sequencer with R9.4.1 flow cells (FLO-MIN106D) from Oxford Nanopore. Adaptive sampling (AS) method was used for the targeted sequencing of regions of interest (ROI). CpGs, including CpG islands, shores, shelves as well as promotor regions, were used as ROI for RRMS. The sequence metrics were updated in **Supplementary Table 5**. Base calling and alignment were processed by guppy/6.2.1_gpu + minimap2/2.26 + remora/2.0.0.

### DiMeLo-seq preparation and analysis

Directed methylation with long-read sequencing (DiMeLo-seq), a method that uses antibody-tethered enzymes to methylate DNA near a target protein’s binding sites in situ, was performed using the previously described protocol with the following modifications[57]. 2 million of D2, D14 and D28 LNCaP+FOXA2 cells or cells from 20mg PDX tumor tissues were used for each DiMeLo-seq. The cells were washed once with 1ml of 1xPBS, then cross-linked with 0.1% formaldehyde for 2 min at room temperature, followed by the addition of 75mM glycine to quench the reaction for 5 min at room temperature. Crosslinked cells were lysed in 1 ml of Dig-Wash buffer (0.02% digitonin, 20 mM HEPES-potassium hydroxide buffer, pH 7.5, 150 mM sodium chloride, 0.5 mM spermidine, 1 Roche cOmplete EDTA-free tablet (11873580001) per 50 ml buffer and 0.1% BSA) for 5 min on ice. The nuclei were bound to Concanavalin A-coated magnetic beads (BangsLaboratoriesBP531) for 10 min at room temperature. The beads-bound nuclei were resuspended in 200 µl of Tween-Wash buffer (0.1% Tween-20, 20 mM HEPES-potassium hydroxide, pH 7.5, 150 mM sodium chloride, 0.5 mM spermidine, 1 Roche cOmplete EDTA-free tablet per 50 ml buffer and 0.1% BSA) containing 4 µl FOXA2 antibody (Abcam, Cat#ab108422) or IgG (CST, Cat#66362), incubated on a rotator at 4 °C for overnight. After primary antibody incubation, the beads were washed twice with 1 ml of Tween-Wash buffer, then beads bound nuclei were gently resolved in 100 µl of Tween-Wash containing 200 nM pA– Hia5 and incubated on a rotator at 4 °C for 2h. After pA-Hia5 binding, the beads were washed twice with 1 ml of Tween-Wash buffer, followed by Hia5 methyltransferase reaction in 100 µl of Activation buffer (15 mM Tris, pH 8.0, 15 mM sodium chloride, 60 mM potassium chloride, 1 mM EDTA, pH 8.0, 0.5 mM EGTA, pH 8.0, 0.5 mM spermidine, 0.1% BSA and 800 µM SAM) and incubated at 37 °C for 2h with replenishing of 800 µM SAM during the incubation. After the incubation, the beads bound nuclei were resuspended in 100 µl of clod PBS containing 2 µl of 20mg/ml Proteinase K, 1 µl of 10% SDS and incubated at 55 °C for 1h. The nanopore DNA sequence libraries were prepared as RRMS described above. The sequence metrics were updated in **Supplementary Table 5**. QC was performed by mosdepth/0.3.4 and NanoStat/1.6.0. Aggregate modified base counts for 5mC and 6mA were performed by modbam2bed/0.9.5. Bigwig file of IGV track was generated by bedGraphToBigWig from kentUtils/v302.1. A and CpG methylation profiles, intensity plots, heatmaps and single DNA molecule tracks were generated by dimelo/0.1.0 python package. Differentially methylated regions (DMRs) were identified by R Bioconductor package DSS/2.48.0. Top 200 hyper and top 200 hypo methylated DMRs nearby genes were annotated by HOMER/4.8.3 and generated in PCA using the Z-score of RPKM value of RNA-seq data.

### Statistical analysis

For each independent *in vitro* experiment, at least three technical replicates were used. Most in vitro experiments were repeated independently three times, and some were repeated twice. Two-tailed unpaired Student’s *t*-tests were used to assess statistical significance in RT–qPCR experiments and cell-based functional assays, and *in vivo* mouse experiments.

### Murine xenograft studies

Mice were housed in specific pathogen-free animal facilities (at 20–23 °C, with 40–60% humidity and 12-h light/12-h dark cycle). LuCaP PDX were derived from resected metastatic PCa with the informed consent of patient donors as described previously [78] under a protocol approved by the University of Washington Human Subjects Division institutional review board. NOD SCID male mice at 6–7 weeks old were purchased from Charles River Laboratories. For LuNE xenograft, 2x10^6 cells were subcutaneously injected into the right flank of the mice. Tumor volumes were measured twice per week with digital calipers, using the formula: V = L × W^2/2 (V, mm3; L, mm; W, mm). Once the size of tumors reached approximately 70 mm^3, the mice were randomized to receive vehicle (5% DMSO:95% methylcellulose) or CCS1477 (20 mg/Kg) every day by oral gavage for 33 days. For LuCaP 145.2 PDX xenograft, fresh tumors were cut into small pieces (8mm^3) using a scalpel and were subcutaneously implanted into both sides of SCID mice. The size of tumors reached approximately 150mm^3 at 1 month after implantation, then mice were randomized to receive vehicle (5% DMSO:95% methylcellulose) or CCS1477 (20 mg/Kg) every day by oral gavage for 18 days. The tumors and other organs of mice were collected for downstream analyses when they reached the endpoint.

### Immunohistochemistry (IHC) and TMA analysis

IHC was performed using ImmPRESS® Excel Amplified Polymer Kit (Vector Laboratories), according to the manufacturer’s instructions. Briefly, Formalin-fixed paraffin-embedded (FFPE) tissue sections were de-paraffinized and hydrated, followed by antigen retrieval was accomplished by heating at 99–100 °C for 15 min in 1x citrate buffer, pH 6.0 (Sigma, C9999-1000ML). Following antigen retrieval, the tissue sections were incubated with BLOXALL blocking solution to quench endogenous peroxidase activity and then blocked with 2.5% normal horse serum. After blocking, the tissue sections were incubated with primary antibodies, followed by incubating with an amplifier antibody (Goat Anti-Rabbit IgG for rabbit primary antibodies) or directly incubated with ImmPRESS polymer reagent (for mouse primary antibodies). After ImmPRESS polymer reagent incubation, the tissue sections were incubated in ImmPACT DAB EqV working solution until desired stain intensity developed, then the tissue sections were counterstained with hematoxylin, mounted with mounting medium, and imaged with an Olympus microscope.

For human TMA staining, TMAs containing metastatic CRPC specimens were obtained as part of the University of Washington Medical Center Prostate Cancer Donor Program, which is approved by the University of Washington Institutional Review Board. All specimens for IHC were formalin-fixed (decalcified in formic acid for bone specimens), paraffin-embedded, and examined histologically for the presence of a non-necrotic tumor. TMAs were constructed with 1-mm-diameter duplicate cores (n = 126) from CRPC tissues (n = 23 patients) consisting of visceral metastases and bone metastases (n = 63 sites) from patients within 8 h of death. TMAs staining of FOXA2 and NKX2-1 were conducted by Northwestern pathology facility. The information of antibodies for IHC was provided in **Supplementary Table 2**. Images were captured with TissueFax Plus from TissueGnostics, exported to TissueFAX viewer, and analyzed using Photoshop CS4 (Adobe). FOXA2 and NKX2-1 immunostaining was scored blindly by a pathologist using a score of 0 to 3 for intensities of negative, weak, moderate, or intense and multiplied by the percentage of stained cancer cells.

## Data availability

All sequencing data (RNA-seq, ATAC-seq, ChIP–seq, Hi-C, RRMS, DiMeLo-seq and single cell sequencing) generated for the study have been deposited in the Gene Expression Omnibus.

## Code availability

The code for NGS analyses performed in this paper are in the process of being uploaded to Github.

## ACKNOWLEDGEMENTS

This work was partially supported by the Northwestern University Pathology Core Facility, Center for Advanced Microscopy/Nikon Imaging Center, and Robert H. Lurie Comprehensive Cancer Center Support Grant (NCI P30CA060553). NGS was done at the University of Chicago Genomics Facility and Northwestern Sequencing Core, Admera Health, and Florida State University College of Medicine. IP-MS was done at Taplin Mass Spectrometry Facility of Harvard Medical School. We thank Dr. Eva Corey (University of Washington) for the generation and maintenance of the LuCaP PDX models that were partially funded by NIH awards P50CA97186 and PO1CA163227. The CRPC TMA was graciously provided by Dr. Colm Morrissey (University of Washington) through PCBN. We thank Dr. Jiaoti Huang (Duke) University) for sharing their scRNA-seq data from clinical PCa samples. We thank Dr. Amina Zoubeidi (University of British Columbia) for sharing enzalutamide-resistant AR+/PSA− cell line 42D. Funding supports for the work include the NIH/NCI training grant T32CA09560 (to GG), prostate cancer SPORE P50CA180995 (JY, XY), R50CA211271 (to JCZ), R01CA227918 and R01CA257446 (to JY), and Prostate Cancer Foundation 2017CHAL2008 (to JY, JCZ).

## Contributions

J.Y. J.C.Z., and X.L. conceived the project and designed the experiments. J.C.Z., V.K., I.C. J.W., F.Y and J.Y. conducted bioinformatic and statistical analyses. G.G. and W.X. assisted with in vivo mouse experiments. X.L. and W.X. performed H&E and IHC experiments. Q.J. performed LNCaP+FOXA2 D14 and D28 Hi-C experiments. X.L. performed the rest of the experiments. N.A. provided pA-Hia5 and consulted on DiMeLo-seq experiments. M.S., P.J., and V.C. provided valuable comments on the project. X.L. V.K., J.Z., and J.Y. wrote the manuscript.

